# A practical guide to linking brain-wide gene expression and neuroimaging data

**DOI:** 10.1101/380089

**Authors:** Aurina Arnatkevičiūtė, Ben D. Fulcher, Alex Fornito

## Abstract

The recent availability of comprehensive, brain-wide gene expression atlases such as the Allen Human Brain Atlas (AHBA) has opened new opportunities for understanding how spatial variations on the molecular scale relate to the macroscopic neuroimaging phenotypes. A rapidly growing body of literature is demonstrating relationships between gene expression and diverse properties of brain structure and function, but approaches for combining expression atlas data with neuroimaging are highly inconsistent, with substantial variations in how the expression data are processed. The degree to which these methodological variations affect findings is unclear. Here, we outline a seven-step analysis pipeline for relating brain-wide transcriptomic and neuroimaging data and compare how different processing choices influence the resulting data. We suggest that studies using AHBA should work towards a unified data processing pipeline to ensure consistent and reproducible results in this burgeoning field.

## Introduction

Over the past two decades, human imaging genetics has emerged as a powerful strategy for understanding the molecular basis of macroscopic neural phenotypes measured across the entire brain (Meyer-Lindenberg and Weinberger, 2006, Muñoz et al., 2009, Arslan, 2015, Hashimoto et al., 2015). Traditionally, this work has involved correlating allelic variation at one or more genetic loci with variation in one or more imaging-derived phenotypes (IDPs), initially through candidate gene studies and more recently at a genome-wide level. The latter has been facilitated by the formation of large consortia, such as ENIGMA (Thompson et al., 2014). A common assumption in this work is that variants associated with an IDP (or nearby variants tagged by the associated variant) influence gene expression or protein abundance, which in turn alters cellular function and ultimately affects the studied IDP. However, multiple environmental and other factors can impact gene activity (Fraser et al., 2005, Choi and Kim, 2007, Cole, 2009) and the functional roles of many IDP-linked variants, which are usually identified through large-scale statistical analyses, are often unknown. As a result, the mechanisms through which a given variant may influence phenotypic variation can be unclear. Moreover, the expression levels of many genes vary substantially across brain regions (Hawrylycz et al., 2015), and these spatial variations cannot be inferred from DNA sequence alone.

Assays of gene expression provide a more direct measure of gene function. Expression assays are invasive, requiring direct access to neural tissue, and technical limitations have historically constrained analyses of gene expression in the brain to small sets of areas studied in isolation. Recent advances in the development of high-throughput tissue processing and bioinformatics pipelines have overcome these limitations, resulting in datasets of gene expression across a large fraction of the genome in a large number of brain regions, and through various stages of development [see Keil et al. (2018) for a detailed overview]. While some of the human atlases span multiple brain areas, only the Allen Human Brain Atlas (AHBA) offers high resolution coverage of nearly the entire brain, comprising expression measures for more than 20,000 genes taken from 3702 spatially distinct tissue samples. Critically, the samples have been mapped to the stereotaxic space, allowing researchers to directly relate spatial variations in gene expression to spatial variations in IDPs (for more details on the AHBA see supplementary material S1).

This unprecedented capacity to link molecular function to macroscale brain organization has given rise to the nascent field of imaging transcriptomics, which has begun to yield new insights into how regional variations in gene expression relate to functional connectivity within: canonical resting-state networks (Richiardi et al., 2015, Forest et al., 2017); fiber tract connectivity between brain regions (Goel et al., 2014); temporal and topological properties of large-scale brain functional networks (Cioli et al., 2014, Vértes et al., 2016); the specialization of cortical and subcrotical areas (Krienen et al., 2016, Parkes et al., 2017, Anderson et al., 2018); regional maturation during embryonic and adolescent brain development (Kirsch and Chechik, 2016, Whitaker et al., 2016); and pathological changes in brain disorders (Rittman et al., 2016, Romme et al., 2017, McColgan et al., 2018, Romero-Garcia et al., 2018a). Software toolboxes to facilitate the integration of brain-wide transcriptomic and imaging data have also been developed (French and Paus, 2015, Gorgolewski et al., 2015, Rizzo et al., 2016, Rittman et al., 2017).

Analyses in imaging transcriptomics are often highly multivariate, involving expression measures of around 20,000 genes in each of around 10^2^–10^3^ brain regions, being related to one or more distinct IDPs quantified in each region requiring quite extensive data processing. The impact of data processing choices on the results of neuroimaging analyses is well documented, with strategies for the correction of motion-related and global signal fluctuations in functional MRI being a prime example (Power et al., 2015, 2017, Ciric et al., 2017, Parkes et al., 2018). Comparable scrutiny has not yet been applied to the many processing choices that can affect the analysis of transcriptomic atlases and their relation to IDPs. At the time of writing, more than 30 studies have linked the AHBA gene expression measures to human neuroimaging data. The lack of a standard processing pipeline for gene expression data means that the degree to which the results of this work are robust to different methodological choices remains unclear.

As the field develops, it is important to establish methodological guidelines to ensure consistent and reproducible results, and to support valid interpretation. In this paper, we offer a practical guide to some of the key steps in processing the AHBA gene expression data and examine the potential impact of methodological choices available at each step. We focus on the AHBA, as it is the most spatially comprehensive and widely used gene expression atlas in the field (Hawrylycz et al., 2012).

The paper is organised as follows. We begin by summarizing some basic aspects of how gene expression is quantified, and general characteristics of the AHBA. We then outline several key steps in a basic workflow for relating gene expression measures to imaging data and examine the impact of methodological choices at each step. In the final section we make some recommendations for best practice, and provide directions for further research.

## Measuring gene expression

Gene expression is a process through which genetic information encoded by sequences of DNA is read and used to synthesize a particular gene product, such as a protein or RNA molecule (Szymański and Barciszewski, 2002). The order of amino acids within each gene determines the structure and function of the resulting product, which in turn affects cellular function and drives phenotypic variability. While the DNA of each cell in the organism is identical, different cells and anatomical structures express different phenotypes (e.g., neurons versus lymphocytes) due to differences in gene expression. The process through which a sequence of DNA is expressed is complex, but (for present purposes) can be divided into two main stages: (1) transcription, which occurs when an unwound segment of DNA is read to produce messenger RNA (mRNA); and (2) translation, which occurs when mRNA is used to synthesize proteins (Krebs et al., 2014). Gene expression is commonly approximated by measuring mRNA levels of a particular gene and is thus an index of gene transcriptional activity. Gene transcription is an indirect proxy for protein abundance, which is ultimately determined by gene translation. This distinction is important as several studies have shown that mRNA and protein levels within a tissue can vary significantly (Futcher et al., 1999, Gygi et al., 1999, Greenbaum et al., 2003) and gene expression (transcriptional activity) and protein abundance (translational activity) are not always positively correlated (Margineantu et al., 2007, Schwanhäusser et al., 2013).

In the AHBA, transcriptional activity has been measured using microarray, which quantifies the expression levels of thousands of genes at once by measuring the hybridization of cRNA (Cy3-labeled RNA) in a tissue sample to a particular spot on the microarray chip. Each of these spots, called probes, maps to a unique location of the DNA and contains single-stranded nucleic acid profiles that are ready to anneal to their complementary targets in the process of hybridization. Relative levels of gene expression in a tissue sample are then quantified by measuring the fluorescence at each sequence-specific location, which is proportional to the amount of complementary mRNA in a sample (Tarca et al., 2006). This method provides a cost-effective way to measure gene expression in high-throughput manner. However, it is limited to known gene sequences, is prone to background noise due to indirect assessment of expression values, and spatial biases can result from variability in lateral diffusion of target molecules on the chip (Steger et al., 2011). Expression measures can also be affected by cross-hybridization artefacts arising when cRNA anneals to an imperfectly matched probe.

Microarray is typically performed on bulk tissue samples, and the cellular composition of a sample can strongly influence its gene expression profile. As a result, two samples with varying densities of different cell types may show transcriptional differences simply because of their different cellular composition. This is an important consideration when comparing data acquired from samples taken from different parts of the brain, since variations in the density of distinct cell types may drive differences in regional gene expression. In addition, variations in the way tissue samples are acquired, handled and processed, age at death (Glass et al., 2013), sex (Berchtold et al., 2008, Trabzuni et al., 2013), ethnicity (Spielman et al., 2007), brain pH (Mexal et al., 2006), post-mortem interval (Zhu et al., 2017), and RNA degradation (Jaksik et al., 2015), can all affect expression measures. Another potential influence arises from batch effects, caused by samples being processed at different times, by different staff, or in different labs; even changing atmospheric ozone levels can impact the final measures (Fare et al., 2003) [see Scherer (2009) for an overview]. The Allen Institute has implemented a series of steps to mitigate this variability as much as possible, as outlined in Allen Human Brain Atlas technical white paper (Allen Human Brain Atlas, 2013).

One final consideration is that any individual gene expression assay provides a static snapshot of a dynamic process. Gene expression changes through development, and as a function of experience, environmental exposures and other factors (Fraser et al., 2005, Choi and Kim, 2007, Berchtold et al., 2008, Cole, 2009, Birdsill et al., 2011, Naumova et al., 2012, Kumar et al., 2013). The further advancement of developmental atlases of gene expression (Johnson et al., 2009, Colantuoni et al., 2011, Kang et al., 2011, Fertuzinhos et al., 2014, Bakken et al., 2016) will help to shed light on these dynamic processes.

### A general workflow for processing brain-wide transcriptomic data

The AHBA consists of microarray data in 3702 spatially distinct samples taken from six neurotypical adult brains. The samples are distributed across cortical, subcortical, brainstem and cerebellar regions in each brain, and quantify the expression levels of more than 20,000 genes (for more details see supplementary material S1). Different brain regions were sampled across each of the six AHBA donors to maximize spatial coverage. Figure 1 shows the variability of coverage across individual brains.

**Fig 1.**
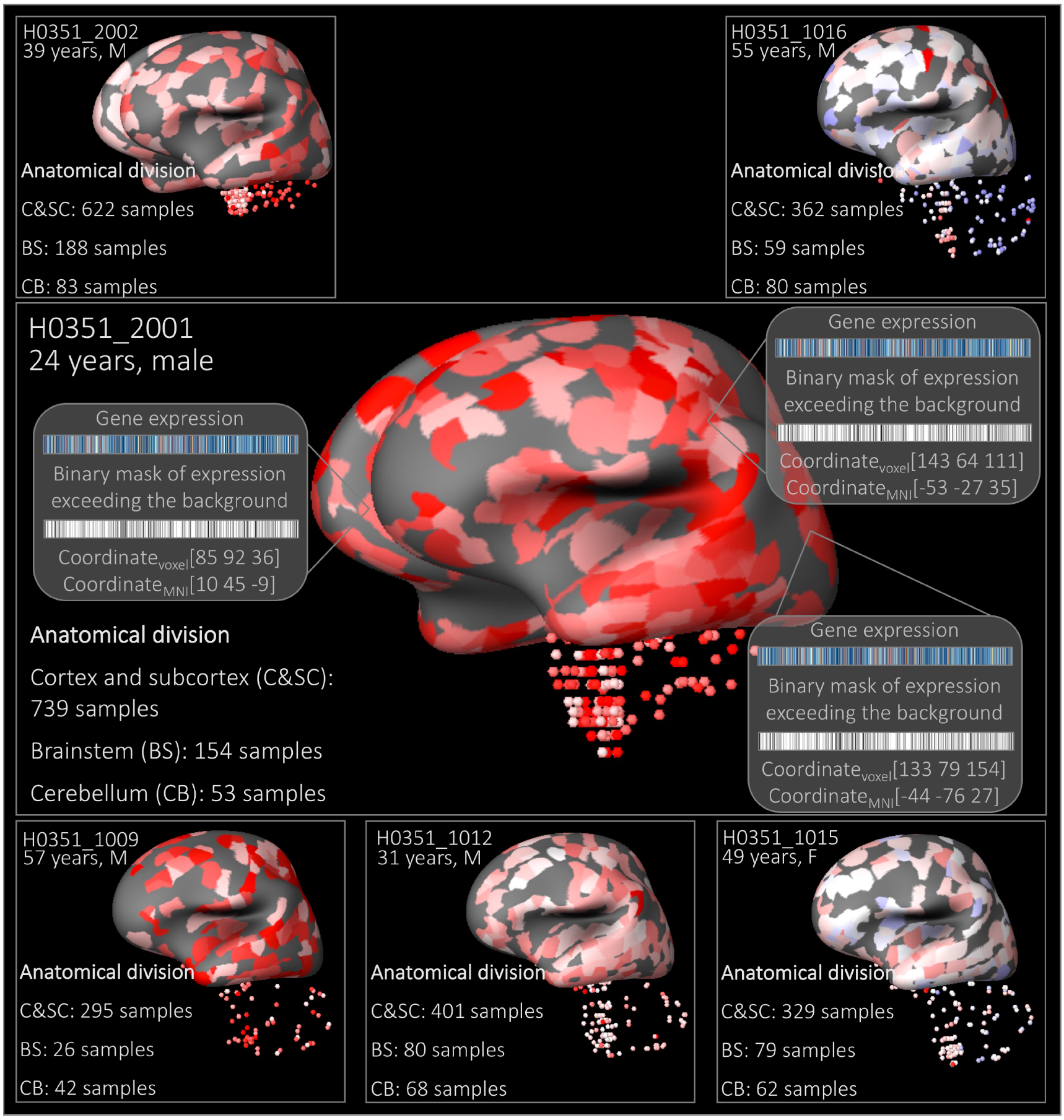
A schematic representation of gene expression data for the CLRN1 gene. (selected for visualization purposes). Discrete tissue samples in each brain are represented as colored areas on a grey brain surface. The color corresponds to the relative gene expression level across all six brains (*z*-score): red (high), blue (low). The size of the patch is not representative of the size of the actual tissue sample used to quantify gene expression which, in reality, was much smaller. The patch has been padded out by the AHBA online platform as a visual aid. The number of samples in each anatomical division: cortex and subcortex (C&SC), brainstem (BS) and cerebellum (CB) for every donor brain is listed. Middle: A schematic representation of available data for each sample which includes expression values for ~20,000 genes, a binary indication of whether the expression levels exceed background noise, and sample native voxel and MNI coordinates for each sample. Brain representation produced using Brain Explorer 2 (2006-2015 Allen Institute for Brain Science. Brain Explorer 2. Available from: http://human.brain-map.org/static/brainexplorer).

Each tissue sample is associated with a numeric structure ID, name and structure label (cortex, cerebellum, or brainstem) in addition to the MRI voxel coordinates in native image space and MNI stereotaxic coordinates, which can be used to match samples to other imaging data (Figure 1). The AHBA also provides: (1) a binary indicator of when the level of a given transcript exceeds background levels, which can be used for quality control purposes; (2) RNA-seq data for a subset of tissue samples in two donor brains (120 samples each), which can be used for cross-validating expression measures (as we show below); and (3) magnetic resonance images, including T1-weighted, T2-weighted, T2-weighted gradient echo and FLAIR scans for all six brains, and diffusion-weighted images for two brains. These scans were collected prior to the dissection for anatomical visualization.

The AHBA samples were processed over approximately three years, which raises concerns about possible batch effects. Expression data were subjected to normalization procedures within a single brain, as well as between brains, to minimize the effect of non-biological biases such as array-specific differences, dissection method, and RNA quality differences among others, while maintaining biologically-relevant variance. Detailed information about the normalization procedures is provided in the technical white paper (Allen Human Brain Atlas, 2013). Despite these procedures, we show below that large inter-individual differences in gene expression remain, such that samples from the same brain tend to have more similar gene expression compared to the samples from other brains. These differences must be taken into account when combining data across all six brains.

Beyond the processing steps applied by the Allen Institute, a number of other steps are required to link expression measures and neuroimaging data. Here we outline seven major steps, which represent the core features of a typical workflow. The data processing steps, summarized in Figure 2, are: (i) verifying probe-to-gene annotations; (ii) filtering of probes that do not exceed background noise; (iii) probe selection, where representative probes (or a summary measure) are selected to index expression for a gene; (iv) sample assignment, where tissue samples from the AHBA are mapped to specific brain regions in an imaging dataset; (v) normalization of expression measures to account for inter-individual differences and outlying values; (vi) gene-set filtering, to remove genes that are inconsistently expressed across six brains and/or to select genes in a hypothesis-driven way based on the research question. (vii) accounting for the spatial patterns in gene expression. The first six processing steps produce the region × gene matrix that can be used for the regional analyses. The final step of accounting for the autocorrelation in the gene expression measures depends on the particular research question. The potential need to account for spatial effects arises because gene expression is more strongly correlated between samples that are separated by short distances compared to those that are far apart, a pattern that has been described in humans (Richiardi et al., 2015, Krienen et al., 2016, Vértes et al., 2016, Pantazatos and Li, 2017), mouse (Fulcher and Fornito, 2016) and *C.elegans* (Arnatkevičiūtė et al., 2018). Although this spatial autocorrelation is, in itself, an important neurobiological feature of the brain transcriptome (Gryglewski et al., 2018), it is critical for any analysis claiming a specific association between spatial variations in gene expression and a given IDP to show that the association exceeds what would be predicted by lower-order spatial gradients of gene expression. In the following sections, we outline some of the choices that can be made at each of these steps and consider their impact on analysis with some recommendations summarized in the conclusions section. Code and data used for data processing and the following analyses are available at github https://github.com/BMHLab/AHBAprocessing and figshare https://figshare.com/sZ441295fe494375aa0c13 respectively.

**Fig 2.**
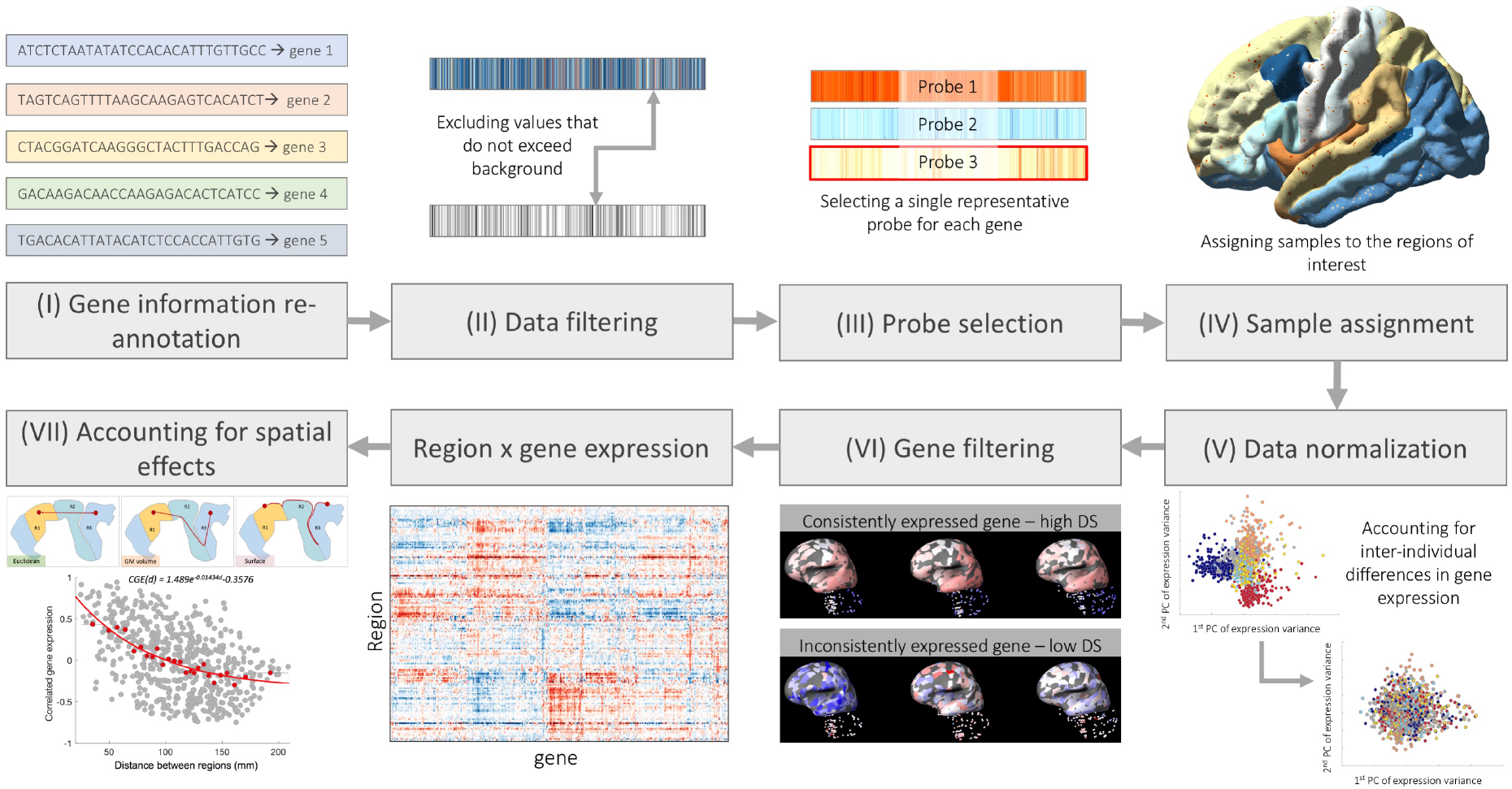
Schematic of a general workflow for processing AHBA to be used in combination with neuroimaging data. The basic workflow involves: (i) confirming and updating probe-to-gene annotations using the latest available data; (ii) background filtering, where expression values that do not exceed background are removed; (iii) probe selection, which, for genes indexed by multiple probes, involves selecting a single representative measure to represent the expression of that gene; (iv) sample assignment, where tissue samples from the AHBA are mapped to specific brain regions in an imaging dataset; (v) normalization of expression measures to account for inter-individual differences and outlying values; (vi) gene-set filtering, to remove genes that are inconsistently expressed across six brains and/or to select genes in a hypothesis-driven way (here we show a gene with consistent expression across three individual brains in the top row and a gene with low consistency in the bottom row, where consistency is measured using a metric called differential stability, or DS (Hawrylycz et al., 2015)). The application of these six steps results in a region × gene expression data matrix that can be further used for the analysis. An important consideration for analyses involving transcriptional data is step (vii) accounting for spatial autocorrelation.

### Step 1. Probe-to-gene re-annotation

In microarray experiments, probe sequences correspond to a unique portion of DNA and are assigned to genes based on available genome sequencing databases (O’Leary et al., 2016). While the AHBA (and other platforms) provide annotation tables where probes are mapped to genes, this information gets outdated with each update of the sequencing databases. For example, at the time of the AHBA release in 2013,18% of probes were not annotated to any gene. Using updated sequencing information we can find corresponding genes for more than 2000 probes that previously were not matched to any gene while some probes are being matched to different genes than before. At the same time some probes can not be unambiguously mapped to any gene using updated sequencing data and therefore should be excluded from further analyses. An accurate probe-to-gene mapping is essential for obtaining biologically meaningful findings. It is therefore necessary to re-assign probes to genes using the most current information available. This re-annotation can be done using several methods and toolboxes, some of which are summarized in Table 1. To our knowledge, only three studies using the AHBA have performed probe-to-gene re-annotation (Richiardi et al., 2015, Eising et al., 2016, Romero-Garcia et al., 2018b).

**Table 1.**
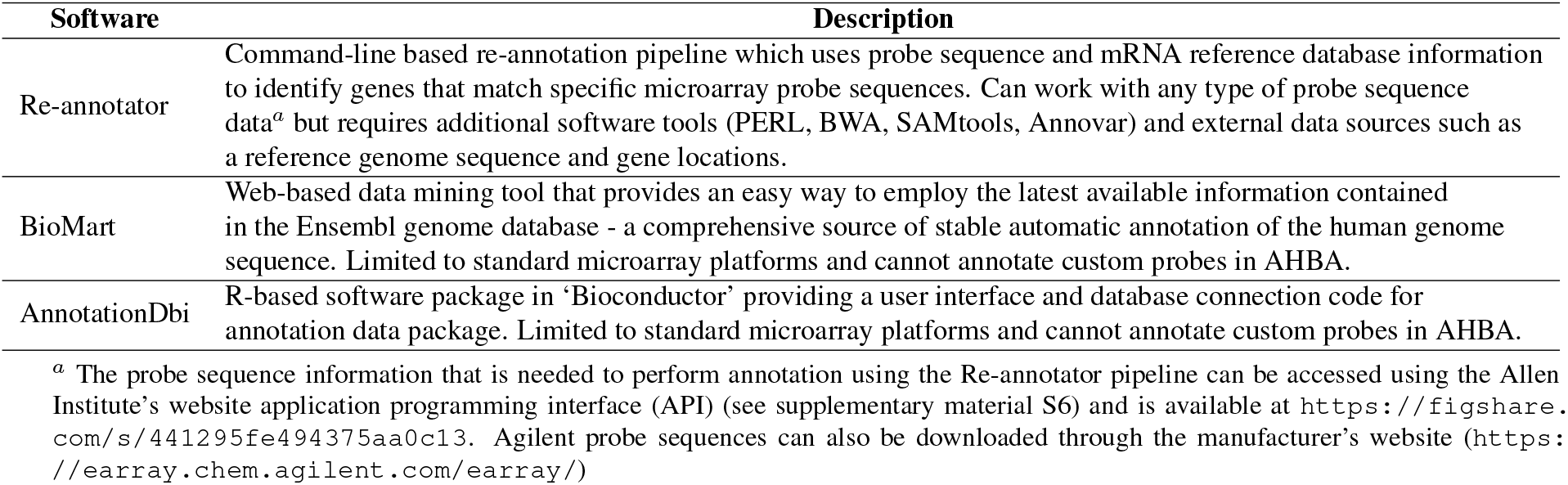
Tools that can be used to update probe-to-gene assignment.

| Software | Description |
| --- | --- |
| Re-annotator | Command-line based re-annotation pipeline which uses probe sequence and mRNA reference database information to identify genes that match specific microarray probe sequences. Can work with any type of probe sequence data<sup>a</sup> but requires additional software tools (PERL, BWA, SAMtools, Annovar) and external data sources such as a reference genome sequence and gene locations. |
| BioMart | Web-based data mining tool that provides an easy way to employ the latest available information contained in the Ensembl genome database - a comprehensive source of stable automatic annotation of the human genome sequence. Limited to standard microarray platforms and cannot annotate custom probes in AHBA. |
| AnnotationDbi | R-based software package in 'Bioconductor' providing a user interface and database connection code for annotation data package. Limited to standard microarray platforms and cannot annotate custom probes in AHBA. |

<sup>a</sup> The probe sequence information that is needed to perform annotation using the Re-annotator pipeline can be accessed using the Allen Institute's website application programming interface (API) (see supplementary material S6) and is available at https://figshare.com/s/441295fe494375aa0c13. Agilent probe sequences can also be downloaded through the manufacturer's website (https://earray.chem.agilent.com/earray/)

To investigate how probe-to-gene annotations change over time, we supplied a list of all available 60 bp length AHBA probe sequences (n = 58,692) to the Re-annotator toolkit (Arloth et al., 2015) (Table 1). We found that 45,821 probes (78%) were uniquely annotated to a gene and could be related to an entrez ID - a stable identifier for a gene generated by the Entrez Gene database at the National Center for Biotechnology Information (NCBI). A total of 19% of probes were not mapped to a gene, and just under 3% were mapped to multiple genes and could not be unambiguously annotated. Of the probes that were unambiguously annotated to a gene, 3438 (7.5%) of the annotations differed from those provided by the AHBA: 1287 probes were re-annotated to new genes and 2151 probes that were not previously assigned to any gene in the AHBA could now be annotated. Additionally, 6211 (~ 10%) probes in the initial AHBA dataset had an inconsistent gene symbol, ID or gene name information according to the NCBI database (https://www.ncbi.nlm.nih.gov/), as of 5th March 2018. Because of these differences, we recommend obtaining probe-to-gene annotations and retrieving the gene symbol ID and name from the latest version of NCBI (ftp://ftp.ncbi.nlm.nih.gov/gene/DATA/GENE_INFO/Mammalia/). Hereafter, we present all analyses using this newly re-annotated set of 45,821 probes, corresponding to 20,232 unique genes.

### Step 2. Data filtering

Microarray experiments are prone to background noise due to non-specific hybridization, so appropriate controls must be employed to discriminate expression signal from noise. Variability in measured intensity values is greater for lower hybridization intensities, where signal levels approach background (Quackenbush, 2002). This problem is often addressed by removing a fixed percentage of probes with lowest intensity or using only array elements that show statistically significant expression differences (increase) from the background (Quackenbush, 2002). Each probe in each sample of the AHBA has been assigned a binary indicator for whether it measures an expression signal that exceeds background levels (see Figure 1). This assignment is done on the basis of two criteria: 1) a *t*-test comparing the mean signal of a probe to the background (*p* < 0.01) indicating that the mean signal of the probe’s expression is significantly different from the background; and 2) the difference between the background and the background subtracted signal is significant (> 2.6 × background standard deviation).

Filtering genes based on the AHBA binary indicator [intensity based filtering (IBF)] can have a marked effect on the final set of genes included for analysis, however only a few published studies using the AHBA data have reported using the IBF (Hawrylycz et al., 2012, Richiardi et al., 2015, Burt et al., 2017). For example, if we exclude probes that did not exceed the background in at least 50% of all cortical and subcortical samples across all subjects, we exclude 30% of probes (13,844 out of 45,821), assaying 4486 out of 20,232 genes (Figure 3A). In other words, if no filtering is performed, > 22% of genes will have expression levels consistent with background noise in at least half of the tissue samples.

**Fig 3.**
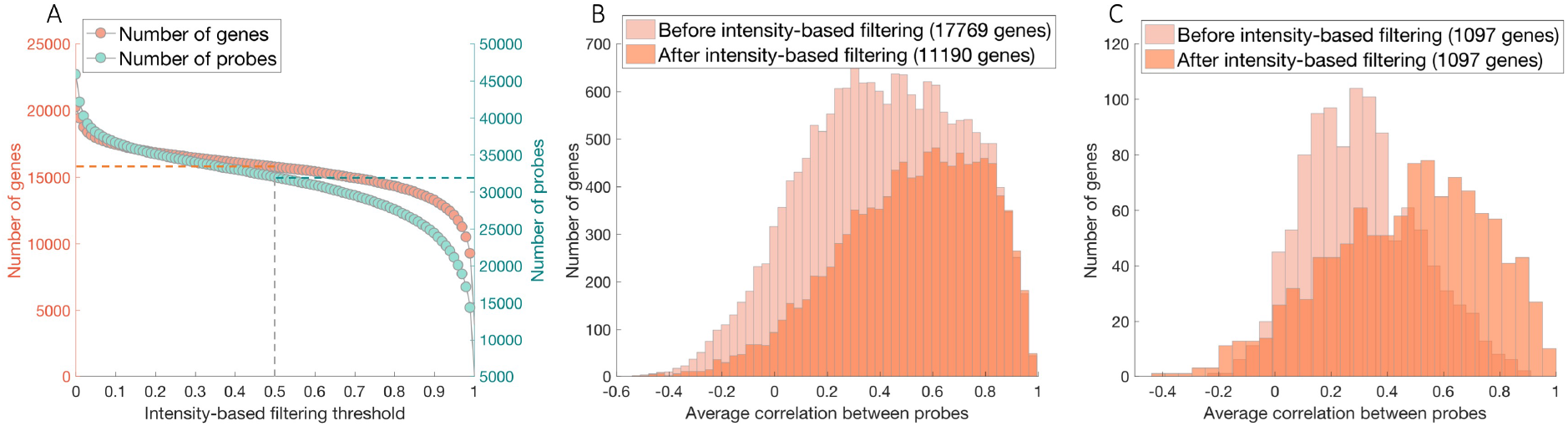
The influence of the intensity based filtering. A) The number of probes and genes as a function of filtering threshold: x axis - the minimum proportion of samples with expression values exceeding the background; y axis - the number of probes and genes retained. Dotted lines correspond to the number of probes and genes retained after 50% filtering threshold was applied. B) Average correlation between expression values measured using all available probes for the same gene: light orange — original set of 17,769 genes with more than 1 probe; dark orange - 11,190 gene set after intensity-based filtering with more than one probe, where probes for which 50% of samples do not survive IBF are removed. C) Distributions of average correlations between expression values measured using all available probes for the same gene that demonstrated any change after IBF (1097 genes, or ~ 10% genes with multiple probes).

To further investigate the impact of IBF, we examined how filtering affects the average correlation between expression values quantified by multiple probes for the same gene. Given that expression measures of different probes are expected to be comparable, IBF should increase the inter-probe agreement. Figure 3B shows the distribution of the average between-probe correlation, estimated before and after IBF. Starting with an initial set of 17,769 genes with multiple probes, applying IBF to exclude probes that do not exceed background in at least 50% of regions removes 6579 genes. It is evident that the distribution of between-probe correlations is pushed towards higher values.

We next compared the mean between-probe correlations obtained before and after IBF, focusing on the 11,190 genes with multiple probes that were retained after filtering. For 10,111 of these genes, the average correlation was identical, while for the remaining 1097 genes (~ 10%), the mean correlation was significantly greater following IBF (Spearman’s rank correlation (denoted as *ρ* through the text): *ρ* = 0.47 vs *ρ* = 0.30; *p* = 7 × 10^−54^, Wilkoxson rank sum test; Figure 3C). Gene score resampling (GSR) analysis (Gillis et al., 2010) revealed that IBF excluded genes that are involved in generic cellular, immunological and metabolic processes that are not specific to the brain (see supplementary file enrichmentExpression.csv for results and supplementary material S2 for more details). While the exact threshold for the IBF still remains to be chosen by the researchers, these results indicate that IBF is effective in mitigating noise in the microarray gene expression measures.

### Step 3. Probe selection

Multiple probes can be used to measure the expression level of a single gene at different exons (segments of RNA molecules that code for a protein or peptide sequence), which can increase the reliability of the measurement. After performing re-annotation and IBF, 71% genes in the AHBA were measured with at least two probes (compared to 93% in the original data). One might expect that probes measuring the expression of the same gene should show consistent expression patterns, but this is not always the case. For example, even after IBF, the correlation between probes measuring the expression levels of the same gene for more than 20% of genes is *ρ* < 0.3 (Spearman rank correlation) (Figure 3B). Investigators have used different strategies to derive a representative measure of gene expression. Some of the strategies used in published work are summarized in Table 2.

**Table 2.** Methods used for deriving an estimate of gene expression in cases where multiple probes are available for the same gene.

| Method | Description |
| --- | --- |
| Mean | Calculate the mean of all available probes for a gene.<br>Tan et al. (2013), French and Paus (2015), Eising et al. (2016), Krienen et al. (2016)<br>Vértes et al. (2016), Whitaker et al. (2016), Negi and Guda (2017), McColgan et al. (2018) |
| Max intensity | Select probe with the highest expression level. Romero-Garcia et al. (2018b) |
| Variance | Select the probe with the highest variance across brain regions. Negi and Guda (2017) |
| PC | Select the probe with the highest loading onto the first principal component of probes. Parkes et al. (2017) |
| DS | Select the probe with most consistent pattern of regional variations across the six donor brains, as quantified using a measure called Differential Stability (DS).<br>Hawrylycz et al. (2015), Kirsch and Chechik (2016) |
| Connectivity variance/intensity | Select the probe with the highest average correlation to other probes<sup>a</sup> for >2 probes;<br>select the probe with maximum variance (connectivity variance) or maximum intensity (connectivity intensity) when only 2 probes available. Hawrylycz et al. (2012, 2015), Myers et al. (2015)<br>Burt et al. (2017), Forest et al. (2017), Anderson et al. (2018) |
| Sequence mismatches | Select the probe with fewest sequence mismatches, or the probe with highest standard deviation if >1 probe have the same number of mismatches<sup>b</sup>. Richiardi et al. (2015) |

<sup>a</sup> This measure is implemented in the R function ‘collapseRows’ from WGCNA package (Miller et al., 2011). Here we used a custom implementation of this function that was applied for samples within each brain separately in order to account for potential inter-individual differences in gene expression (discussed in step 5).
<sup>b</sup> Sequence mismatch is defined by Re-annotator software as the difference between probe sequence and the reference genome sequencing data. Considering that ~ 93% probes were re-annotated without any mismatches, the overwhelming majority of probes will be chosen applying the highest standard deviation criteria and resemble maximum variance selection approach. Therefore, this probe selection criteria was not implemented separately and is not presented in the probe selection comparison plots.

To evaluate how the gene expression measures vary under different probe selection methods, we estimated, a single summary measure of expression for each gene indexed by multiple probes, according to one of the methods listed in Table 2. We also evaluated a few other methods beyond ones used in the previous literature, such as selecting a probe with maximum coefficient of variation across samples (CV), or the probe with the highest proportion of samples with expression levels exceeding background noise (signal proportion). In addition, we included random probe selection (averaged over 100 repeats) for comparison. We then took the expression vector of each gene across tissue samples and computed the Spearman rank correlation coefficients between these vectors estimated for each possible pair of methods.

Figure 4A shows the average correlation between expression measures selected using different criteria, averaged across 17,769 genes - all genes with multiple probes available for the same gene. Since most studies using the AHBA do not report using IBF, we show results for unfiltered data (similar results have been obtained using data after IBF, see Figure S2). The average correlation coefficients between probe selection methods range between 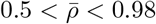, indicating that the probe selection method can have a major impact on expression estimates. The method of summarising the expression measures for a gene as the mean across all available probes is the most highly correlated, on average, to all the other methods. Variance-related methods [coefficient of variation, maximum variance, connectivity-variance and highest loading on first PC of non-normalized data (Parkes et al., 2017)] are similar to each other, but different to other methods. Consistency (DS) and intensity (max intensity, signal proportion and connectivity-intensity) related methods, on the other hand, are more correlated with each other. Notably, the correlations between gene expression measures selected based on the highest CV compared to the consistency/intensity-related criteria are much lower than resulting from the random probe selection strategy, indicating that these methods favour dissimilar properties of expression measures for probe selection. A more detailed discussion of these results is presented in supplementary material S3.

**Fig 4.**
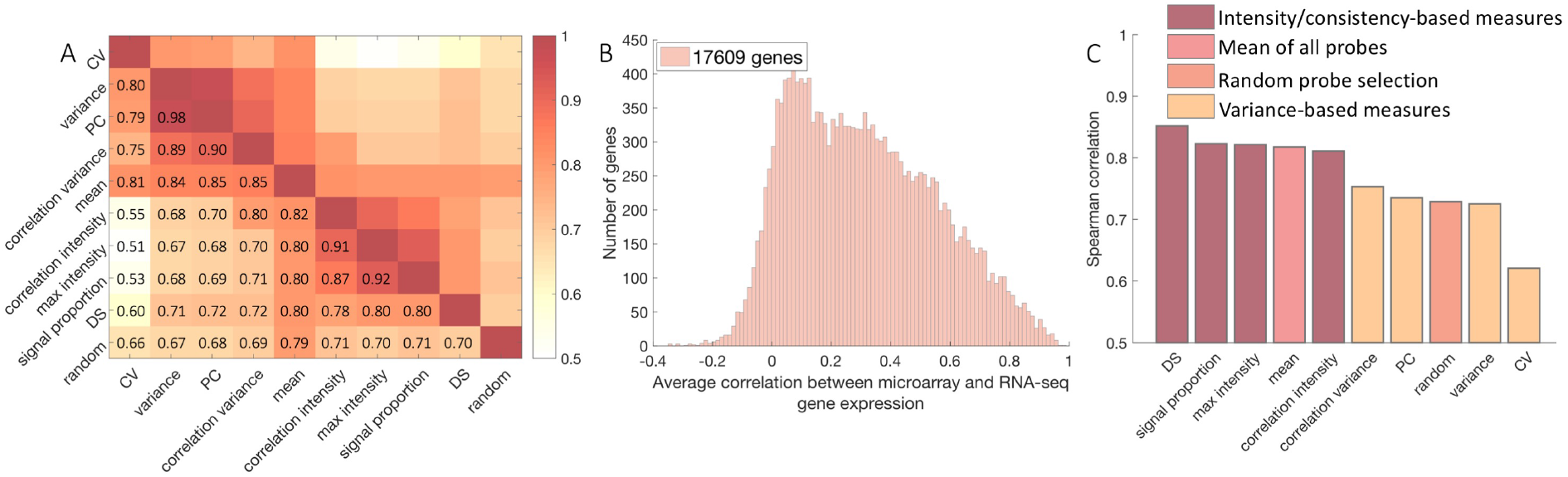
Probe selection method can have a large effect on resulting gene expression estimates. A) Mean Spearman correlation coefficient of expression levels across 17,769 genes with multiple probe annotations, using a range of different probe selection methods: CV, variance, PC, signal proportion, DS, correlation variance, correlation intensity, mean (see Table 2) or selecting a representative probe at random (correlation values averaged over 100 runs). The average correlation is computed over 17,769 genes with multiple probe annotations. B) The distribution Spearman correlation values between microarray and RNA-seq expression data for genes that are present in both datasets. When multiple probes for a gene are available, the maximum correlation value between probes was selected. C) Average correlation between probes selected using RNA-seq (i.e., by selecting the probe that is most correlated with the RNA-seq data) and other methods (ordered by decreasing values, based on 10,221 genes that: (i)had more than one probe available; (ii) were present in both microarray and RNA-seq datasets; (iii) were correlated to RNA-seq (*ρ* > 0.2, Spearman’s rank correlation) to ensure that RNA-seq based probe selection provides a meaningful estimate).

The lack of a gold standard makes it difficult to choose between different probe selection options. One strategy is to use RNA-seq data as an external reference (Miller et al., 2014b). RNA-seq allows precise quantification of the amount of RNA in the sample without reliance on existing knowledge about genome sequences [for an overview, see Wang et al. (2009), Kukurba and Montgomery (2015)]. It is also free of the background noise artefacts that are known to contaminate hybridisation-based gene expression measures and therefore provides a more reliable estimate of gene expression. Samples from two AHBA brains previously analyzed using microarray were reprocessed using RNA-seq to provide expression data for more than 20,000 genes, in 120 samples in each brain (Hawrylycz et al., 2012). Comparing expression values for matching structures in each of the two brains allows us to select probes that correlate most strongly with RNA-seq, providing an additional quality control measure to cross-validate probe selection.

Considering that 17,609 of the 20,232 genes in the microarray data have RNA-seq measures, we first aimed to evaluate whether excluding the ~ 13% of genes that do not overlap between the datasets would eliminate brain-relevant genes. We verified this using over-representation analysis ORA: the genes removed are not enriched in brain-specific functionality but rather are related to septin assembly and organization, as well as the negative regulation of RNA splicing (see supplementary file enrichmentExpression.csv for results and supplementary material S2 for more details).

We then examined the correlations between microarray and RNA-seq expression measures in the 17,609 genes that overlap between both RNA-seq and microarray datasets across 112 brain regions, as shown in Figure 4B. Most correlations are low, with 52% of genes exhibiting a correlation *ρ* < 0.3 and only 23% genes exhibiting a correlation *ρ* > 0.5. This divergence between RNA-seq and microarray is likely to be caused by inaccuracies in the microarray measurements. Using GSR analysis (Gillis et al., 2010) we find that genes with higher correlations between microarray and RNA-seq are related to neuronal connectivity and communication related processes with categories such as ‘transmission of nerve impulse’, ‘ensheathment of neurons’, ‘myelination’ and ‘glial cell development’ demonstrating the strongest enrichment (see supplementary file enrichmentExpression.csv for results and supplementary material S2 text for more details). This analysis demonstrates that RNA-seq data can be used as a reference to select brain-relevant and reliably measured genes.

Figure 4C shows that, compared to other probe selection methods, RNA-seq demonstrates the highest similarity to intensity/consistency-based approaches (*ρ* > 0.8, Spearman’s rank correlation), with DS showing the highest correlation. In contrast, variance-based methods are no more similar to the RNA-seq measures than random probe selection (*ρ* < 0.75, Spearman’s rank correlation). Given that RNA-seq data is only available for a limited number of samples (with only 87% of genes being represented), and the data come from only two of the six brains donor brains in the AHBA, Figure 4C indicates that DS may be a reasonable alternative method for probe selection that can be generalized to the full AHBA.

### Step 4. Assigning samples to brain regions

The AHBA provides gene expression data for multiple spatially localized tissue samples (Figure 1). When relating such data to macroscopic IDPs, it is necessary to generate some mapping between the spatial location of each tissue sample and the particular spatial unit of analysis (e.g., voxel, brain region) used to construct the IDP. This mapping is facilitated by the AHBA including an MNI coordinate (and voxel coordinate) for each tissue sample, and MRI data acquired for each individual brain contained in the AHBA. Each tissue sample is also associated with an anatomical structure ID, which can be related to corresponding higher order structures using the Allen Institute anatomical ontology, allowing brain structures to be identified at different resolution scales.

Existing studies have used several approaches to map tissue samples to regions-of-interest (ROIs) in imaging data. One strategy has been to match samples to structures based on the name of a given anatomical sample. The simplest approach is to use the anatomical structure names provided by the AHBA [see Allen Human Brain Atlas (2013), Tan et al. (2013), Myers et al. (2015), Chen et al. (2016), Kirsch and Chechik (2016), Hecker et al. (2017), Lee et al. (2017), Negi and Guda (2017)], but these regions do not directly correspond to brain parcellations typically used in imaging analyses, so precise alignment with imaging data can be difficult. An alternative approach is to use the MNI (or voxel) coordinates of each sample (Goyal et al., 2014, Cioli et al., 2014, French and Paus, 2015, Richiardi et al., 2015, Komorowski et al., 2016, Krienen et al., 2016, Rizzo et al., 2016, Burt et al., 2017, Parkes et al., 2017, Romme et al., 2017, Shin et al., 2017, Anderson et al., 2018, Romero-Garcia et al., 2018b). It is possible to either assign samples to brain regions in a single parcellation defined in MNI space (Krienen et al., 2016, Keo et al., 2017, Parkes et al., 2017, Romme et al., 2017), or to assign samples to regions based on parcellations of each individual AHBA brain (Romero-Garcia et al., 2018a). The former approach is simpler, but a characteristic of the AHBA is that the MNI coordinates provided for each tissue sample are based on spatial normalizations that were tailored to each individual brain. Specifically, two of the AHBA brains were scanned *in cranio* and normalized to MNI space via a linear transformation, whereas the other four were acquired *ex cranio* and normalized using an affine followed by a non-linear transformation [for details see (Allen Human Brain Atlas, 2013)] with the deformation fields also being smoothed to facilitate matching the images to Nissl stains. These differences across brains will influence the accuracy of the normalization across the six brains, which is compounded by differences in tissue distortion that occurred during sample handling and processing.

To overcome these issues, a parcellation scheme can be applied to each individual donor brain. This method can more accurately account for individual differences in donor brain anatomy but is contingent on being able to generate appropriate transformations between native and MNI space for an accurate parcellation. For cortex, the accuracy of the parcellation can be greatly enhanced by parcellating and normalizing the surface; parcellation of non-cortical areas requires volumetric normalization. In our own work, we have been able to segment the cortical surfaces of the six AHBA brains with reasonable accuracy (assessed by visual inspection), and we supply four different volumetric parcellations mapped at different resolutions to each brain: the Desikan-Killany (Desikan et al., 2006), comprising 34 nodes per hemisphere, the group-level HCPMMP1 (Glasser et al., 2016) comprising 180 nodes per hemisphere, and two random parcellations comprising 100 and 250 nodes per hemisphere, respectively.

Once a particular parcellation has been generated, tissue samples should be assigned to the nearest region of the parcellation. In this assignment, a threshold can also be applied, to avoid assigning samples beyond a certain distance threshold. The distance between sample and region is commonly estimated as the Euclidean distance in 3D space. This sample-region distance has been computed in different ways, including representing a region in space by its centroid coordinate (Vértes et al., 2016, Whitaker et al., 2016, McColgan et al., 2018), or taking the minimum distance between the sample and any voxel in the region (French and Paus, 2015, Parkes et al., 2017, Romme et al., 2017). The latter approach is more accurate, given that regions in any given parcellation vary in size and folded geometry (e.g., Figure 5A).

**Fig 5.**
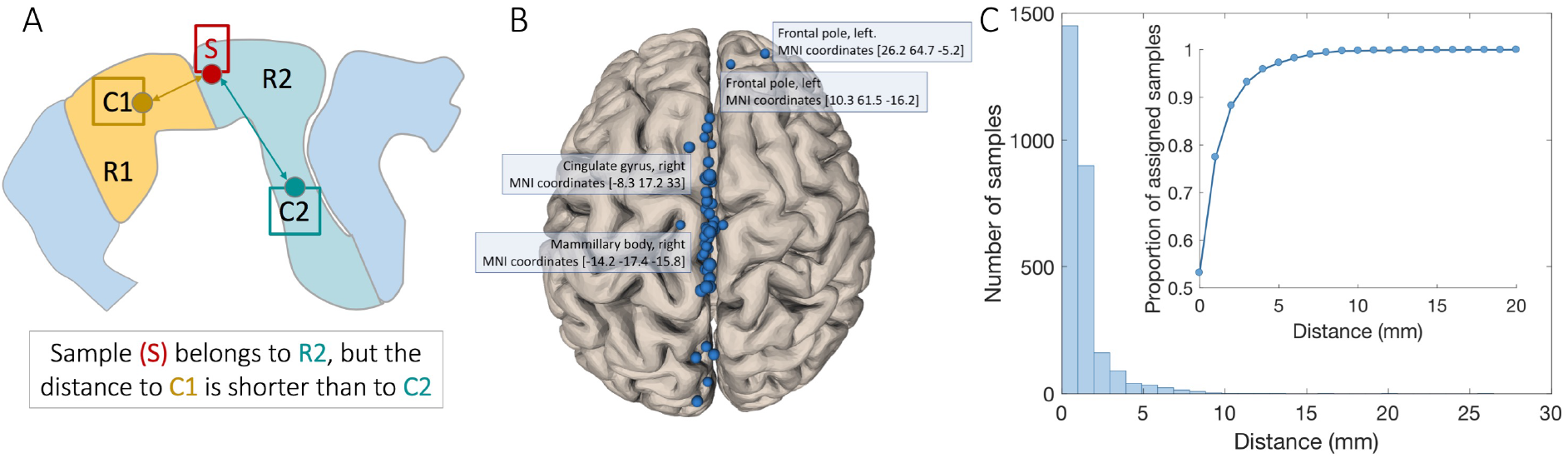
Methods for assigning localized tissue samples to matching regions in a brain parcellation are sensitive to the metric used to define sample-region distances, the distance threshold used, and the use of anatomical annotations on individual samples. A) Schematic representation of sample assignment when a sample is assigned to the closest ROI centroid. A given sample belongs to region R2 but is closer to the centroid (C1) of region R1 than the centroid (C2) of R2, resulting in an erroneous assignment. B) Schematic representation of samples that were assigned to a hemisphere that differed from the annotations provided with their MNI coordinates. C) Sample assignment using distance thresholds: the number of samples across all six brains within a given distance from a parcellation region. Insert shows the proportion of assigned samples as a function of distance threshold. At 2 mm distance threshold, ~90% of tissue samples can be matched to a region in the parcellation.

In this process of assigning samples to regions, errors can occur if the mapping is not done separately for (i) broad anatomical division (cortex, subcortex, cerebellum and so on); and (ii) left and right hemispheres. That is, cortical samples listed as coming from the left hemisphere in the AHBA ontology should only be mapped to left cortical voxels (as samples were taken from annatomically known positions in the brain), right subcortical or cerebellar samples to right subcortical/cerebellar voxels and so on. In our own experience, we have observed that subcortical samples (as indicated by AHBA ontology) can be mapped to cortical regions of the parcellation as cortical voxel may be closer (or visa versa). Similarly, if no separation between hemispheres is performed, 58 out of 2748 cortical and subcortical samples are assigned to an incorrect side of the brain when using the Desikan-Killany (Desikan et al., 2006) parcellation (Figure 5B). While the majority of those samples are very close to the midline, several are clearly incorrectly mapped to the stereotaxic space, such as two samples in the frontal pole, which are assigned to the left side of the brain according to the AHBA annotations but have a positive MNI *x*-coordinate. The same is true for some samples from the mammillary body and cingulate gyrus, which are labelled as coming from the right hemisphere but have negative MNI *x*-coordinates (Figure 5B). To avoid potential mistakes, samples with mismatching assignment should be excluded.

A second consideration is to set a distance threshold for assigning samples to regions, to ensure that samples further than a given threshold away from the parcellation will not be assigned. As shown in Figure 5B, only around 50% of samples are directly mapped to a parcellation when using the Desikan-Killany (Desikan et al., 2006) parcellation (i.e., their coordinates correspond to a voxel inside the parcellation). Increasing the distance threshold will allow some tolerance for small errors in spatial normalization. Figure 5C shows that assigning samples that are up to 2 mm away from any voxel in the parcellation increases the proportion of assigned samples to almost 90%, with additional increases in the distance threshold yielding only minor gains in the number of assigned samples, therefore, we use a 2 mm distance threshold in our analyses.

### Step 5. Six brains, one atlas: accounting for individual variability

In cases where a given brain region is assigned multiple samples, we must generate some aggregate measure of expression for that region. Most commonly this is done by taking a mean across the samples assigned to a given region. A complication of the AHBA is that the samples come from different brains. As we shown in the next section, each brain shows a distinct transcriptomic profile, which must be addressed before data from different brains can be aggregated.

The AHBA is often used to represent a general transcriptomic profile of the adult human brain. However, it is comprised of data taken from people aged 24 to 57 years, of different ethnicities, sexes, medical histories, causes of death, and post-mortem intervals (Table 3). Many of these factors can impact gene expression (Fraser et al., 2005, Berchtold et al., 2008, Kumar et al., 2013, Trabzuni et al., 2013). One way to address this brain-specific variance is to conduct analyses separately in each brain. However, spatial coverage of different brain areas in the AHBA varies from person to person, therefore, collapsing samples from all brains allow to derive a single atlas with maximum spatial coverage across the brain. In this case, an appropriate correction for donor-specific transcriptomic differences is required.

**Table 3.** Additional information about the six adult donors in AHBA.

| Donor | Age | Sex | Ethnicity | Medical conditions | Post-mortem interval<sup>a</sup> |
| --- | --- | --- | --- | --- | --- |
| H0351_2001<sup>b</sup> | 24 | Male | African American | History of asthma | 23 hours |
| H0351_2002<sup>b</sup> | 39 | Male | African American | Possible small pituitary adenoma; single neurofibrillary tangle in entorhinal cortex | 10 hours |
| H0351_1009 | 57 | Male | Caucasian | History of atherosclerotic cardiovascular disease | 25.5 hours |
| H0351_1012 | 31 | Male | Caucasian | Sudden cardiac arrest; benign spindle cell proliferation and dystrophic calcification in temporal horn of lateral ventricle | 17.5 hours |
| H0351_1015 | 49 | Female | Hispanic | Splenectomy, hypothyroidism treated with Levothroid; modest numbers of hemosiderin laden macrophages noted in Virchow-Robin spaces in parietal and occipital lobes, mild arteriosclerosis | 30 hours |
| H0351_1016 | 55 | Male | Caucasian | Coronary artery atherosclerosis, prescriptions for clotting and high cholesterol | 18 hours |

<sup>a</sup> Post-mortem interval is defined as the time period from the time of death to the time the tissue is frozen.
<sup>b</sup> These donors have tissue samples collected across both left and right hemispheres while all the other donors have samples only from within the left hemisphere.

Considering that Allen Institute applied a range of data normalization procedures to remove batch effects and artefactual inter-individual differences, most studies using AHBA have not taken into account the additional interindividual differences that might be important when aggregating data across six donor brains. Here we investigated whether intrinsic inter-individual differences in expression play a major role by projecting all tissue samples from six donor brains into a two-dimensional transcriptional principal components space. Figure 6A plots loadings of each cortical tissue sample on the first two principal components of gene expression for all six donors (for the whole brain see Figure S3). This unsupervised projection of samples into gene expression space captures the latent dimensions of variance between all samples and broadly separates the six donors (regardless of where a tissue is located in the brain), indicating that each donor has a distinctive gene expression profile. In other words, while the data normalization procedures applied by the Allen Institute prior to data release removed batch effects and artefactual inter-individual differences, a considerable degree of intrinsic donor-specific variance remains and must be accounted for in order to perform valid data aggregation.

**Fig 6.**
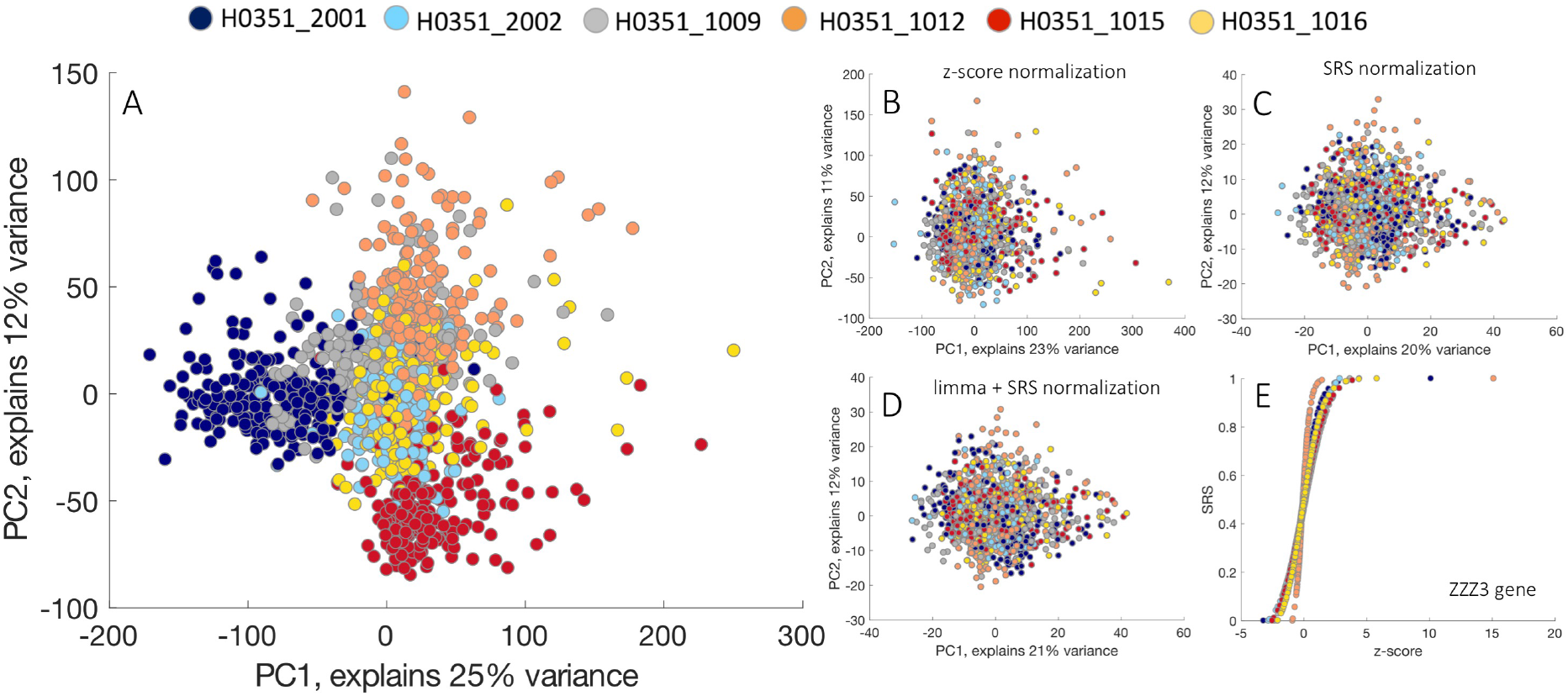
Without applying individual-specific normalization, inter-subject differences dominate cortical gene transcription profiles. A) Non-normalized gene expression data in principal component space. Data from different donors are represented in different colours. Samples from different subjects occupy different parts of the low-dimensional gene expression space. Panels B,C represent gene expression data in principal component space normalized separately for each subject using *z*-scores (B) or the scaled robust sigmoid (SRS) transform (C). Panel D shows gene expression data in principal component space after applying *limma* batch effect removal on cross-subject aggregated data, followed by SRS normalization. After normalization (B,C,D), samples no longer segregate by donor. E) Correlations between normalized expression values (*z*-score vs SRS) for the ZZZ3 gene (chosen for visualization purposes). Each dot represents a normalized expression value for ZZZ3 gene across samples, and different colours correspond to different subjects. The *z*-score normalization results in extreme values being assigned to outliers, therefore producing different scales for subjects and complicating direct comparison of values. SRS normalization is not affected by the outliers and produces normalized values on the same scale for each subject. This example demonstrates how outliers in the data can affect the scaling for different subjects: using *z*-score normalization results in different scales for different subjects with normalized values ranging from approximately −5 to 5 for five out of six subjects, whereas subjects H0351_1012 (orange) and H0351_2001 (dark blue) have a much wider range. In comparison, SRS produces normalized expression values on the same scale for each subject without being affected by outliers. Representations in principal component space are based on 10,028 genes, with representative probes chosen based on correlation to RNA-seq data.

One approach for addressing donor-specific effects is to perform a leave-one-out analysis, where the analysis is repeated six times, excluding one of the brains at each iteration (Parkes et al., 2017, McColgan et al., 2018). This approach can ensure that the results are not driven by single brain. A more direct way of eliminating the inter-individual differences in expression measures is to normalize the gene expression data separately for each subject (Rizzo et al., 2016, Liu et al., 2017, Negi and Guda, 2017, Romme et al., 2017, Romero-Garcia et al., 2018a). With this approach, each gene’s expression values are normalized across regions separately for each donor in order to reflect the relative expression of each gene across regions, within a given brain (Figure 6B-D). A desirable normalization procedure should offer robustness to outlying values and quantify expression on the same scale across donors to enable direct comparison. Most studies using AHBA have generally used *z*-score normalization (Rizzo et al., 2016, Negi and Guda, 2017, Romero-Garcia et al., 2018a),

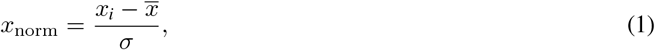

where 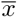 represents the mean, *σ* represents the standard deviation and *x_i_* — the expression value of a gene in a single sample. The estimates of 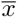 and *σ* are appropriate for symmetric distributions, whereas gene expression distributions across brain samples are often non-symmetric, and can contain outliers, which can bias these summary statistics. Figure 6E demonstrates the sensitivity of *z*-score normalization to the outlying values. A variety of outlier-robust normalizations exist such as Hampel hyperbolic tangent transformation, however here we focus on a variant of a normalization method used by Fulcher and Fornito (2016), the scaled robust sigmoid (SRS) normalization (Fulcher et al., 2013). This approach normalizes gene expression values based on an outlier-robust sigmoid function,

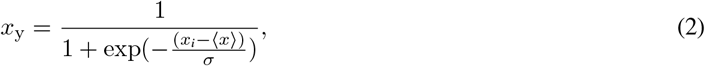

where 〈*x*〉 represents the median and *σ* represents the standard deviation, before rescaling normalized values to a unit interval,

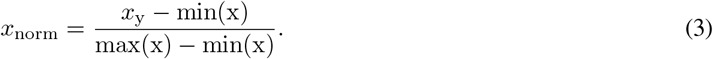

This normalization is robust to outliers and ensures equivalent scaling of expression values for each person. Figures 6C and E show the effectiveness of SRS in dealing with outliers and scaling. Other strategies for removing donor-specific effects involve using linear models applied to cross-donor combined data. For example, donor-specific effects can be treated as an additional batch effect and removed via linear modelling using the R/Bioconductor software package *limma* (Ritchie et al., 2015). While this approach removes inter-individual differences in gene expression, the linear model is sensitive to outliers. This correction in turn can be followed by SRS normalization to minimize the influence of outliers (Figure 6D).

To account for potential between-sample differences in gene expression, Burt et al. (2017) introduced within-sample normalization across genes before subject-specific normalization across samples (see Figure S4). Indeed, some samples can show a markedly different expression profile (extremely low or high values across all genes) from other samples in close spatial proximity that may be caused by measurement artefacts. The influence of these artefacts can be minimized by applying within-sample cross-gene normalization to quantify relative expression levels within a given sample, before normalizing across samples. To quantify the effect of the initial within-sample normalization, we calculated the correlations between expression values across genes and samples in two cases: i) when only cross-sample normalization for each gene was applied; ii) when both cross-gene normalization within sample as well as cross-sample normalization for each gene were applied. While the correlation values were relatively high (median_sample_(*r*) = 0.969, IQR = 0.04; median_gene_(r) = 0.856, IQR = 0.1), the initial within sample normalization was beneficial in reducing potential measurement artefacts in the data.

One additional consideration is that the spatial distribution of tissue samples across individual brains in the AHBA is not uniform. As such, different brains can contribute a different number of samples to any given brain region (Figure 1 and Figure 7). In light of this variability, we have two choices: we can either average all samples falling within a region, meaning that the average may be driven by a subset of individuals who have more samples localized to that region, or we can average at the level of each individual donor brain before aggregating across people (Figure 7). The latter approach ensures that each donor makes an equal contribution to the mean, provided that all genes are normalized to the same scale, however, the choice between those two options can be made depending on the researchers preference.

**Fig 7.**
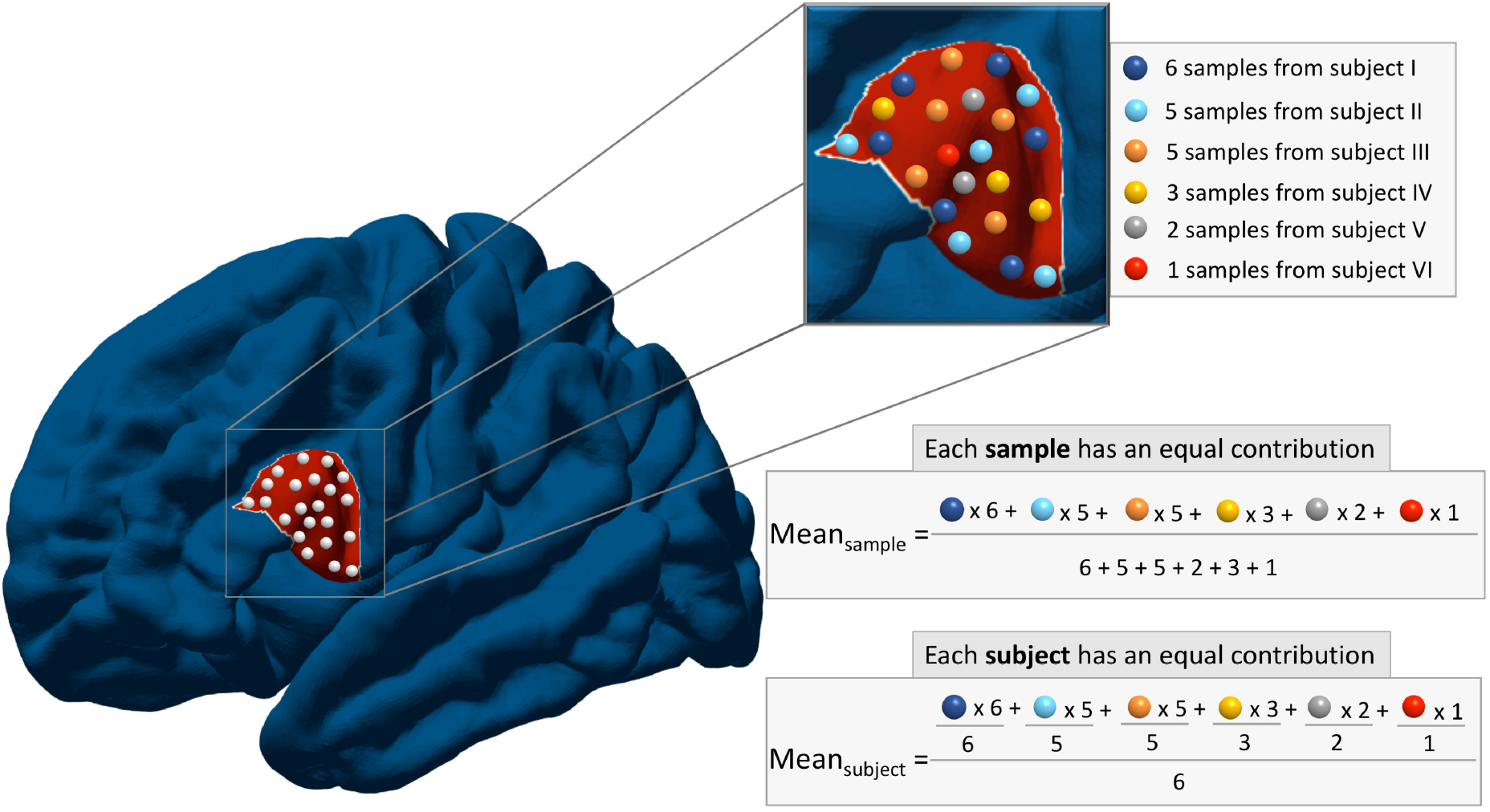
Two alternative ways of averaging expression values within the region of interest. Each region of interest might contain a different number of samples from each subject. For example, the presented region has 6 samples from subject I, 5 samples from subjects II and III, 3 samples from subject IV and only 2 and 1 samples from subjects V and VI respectively. One way of calculating the mean is to take the average of all samples, such that each sample makes an equal contribution to the overall expression value (Mean_sample_). An alternative is to average the samples from each donor first, and then take a second-order average across donors (Mean_subject_). The latter ensures that each subject makes an equal contribution to the summary expression value. The former might be influenced by donors with more samples in a given region.

### Step 6. Gene filtering

The AHBA consists of more than 20,000 unique genes, of which only a fraction is expected to show consistent regional variations in expression across the brain. Many analyses interested in transcriptomic signatures of IDPs will be primarily interested in these brain-specific genes. Various methods for pre-selecting genes of interest have been adopted, including selecting: (i) disease-specific genes (Rittman et al., 2016, Romme et al., 2017, Yokoyama et al., 2017), (ii) genes related to a priori hypotheses (Goyal et al., 2014, Komorowski et al., 2016, Krienen et al., 2016, Acevedo-Triana et al., 2017), or (iii) genes that are expressed consistently across all six AHBA brains, as quantified using the DS measure (Hawrylycz et al., 2015). Genes with high DS values demonstrate consistent patterns of regional variation in expression across the six AHBA subjects, and have been shown to be enriched for brain-related biological function (Hawrylycz et al., 2015). Filtering based on DS thus offers a more targeted approach for investigating relationships between IDPs and gene expression compared to the whole-genome analysis. The selection of disease-specific genes is traditionally based on previous GWAS studies (Satake et al., 2009, Simón-Sánchez et al., 2009, Höglinger et al., 2011, Ferrari et al., 2014, Ripke et al., 2014, Kouri et al., 2015), while gene selection based on an *a priori* hypothesis can depend on other factors such as a specific involvement in clinical disorders (Komorowski et al., 2016, Acevedo-Triana et al., 2017). One particular set of 19 genes demonstrating a selective enrichment in the upper layers of the human cortex compared to mouse [Human Supragranular Enriched (HSE) genes] has been extensively investigated and was found to be implicated in both the functional (Krienen et al., 2016) and topological organisation of the brain (Vértes et al., 2016, Romero-Garcia et al., 2018b). While selecting an appropriate gene filtering strategy rather than implementing the analyses on the whole-genome data is a highly research question-specific choice, investigating the relationships between IDPs and the patterns of gene expression using AHBA may benefit from the initial DS-based filtering.

### Step 7. Accounting for spatial effects

The application of steps 1 to 6 results in a processed region × gene matrix of transcription level values, which can be used for further analyses. Typically, the data are linked to IDPs at either the regional level, or at the level of pairs of regions (i.e., patterns of correlated gene expression, or CGE, between pairs of brain regions are related to pair-wise measures of structural or functional connectivity between those regions). In both cases, we seek to understand how spatial variations in gene expression or CGE relate to spatial variations in the IDP. One complicating factor is that cortical regions that are located in close proximity are more likely to share similar gene expression patterns (Richiardi et al., 2015, Krienen et al., 2016, Vértes et al., 2016, Pantazatos and Li, 2017, Richiardi et al., 2017). A similar spatial autocorrelation of gene expression has been reported in the mouse brain (Fulcher and Fornito, 2016) and in the head of the nematode *C. elegans* (Arnatkevičiūtė et al., 2018). In some respects, this is an interesting and physiologically meaningful trend that warrants further investigation. However, if an IDP varies across the brain in a manner that reproduces a spatial gradient in gene expression, any apparent association between the IDP and gene expression measures may be driven by low-order spatial effects. Depending on the research question, especially when a direct relationship between an IDP and gene expression is evaluated, it is important to confirm that the identified association is stronger than what would be expected based on the spatial autocorrelation properties of gene expression (if such an effect is claimed).

A critical first step in understanding spatial biases in gene expression is to define distances between brain regions. These distances can be estimated by (i) calculating the Euclidean distance between regions; (ii) estimating the shortest distance within the grey matter volume; or (iii) estimating the shortest distance on the cortical surface (Figure 8), see supplementary material S4 for more details. The Euclidean distance is the simplest method, but it approximates distances as straight lines that do not respect cortical geometry. Calculating distances within the grey matter volume or on the cortical surface present a more biologically reliable approach, as distances are quantified considering the cortical geometry. A comparison of these methods, shown in Figure 8D, demonstrates that evaluating the Euclidean distance results in shorter distances, on average, compared to other methods, while anatomically constrained volume and surface-based approaches yield similar distance estimates in cortex. Note that only the Euclidean approach can be generalized for measuring distances to subcortex.

**Fig 8.**
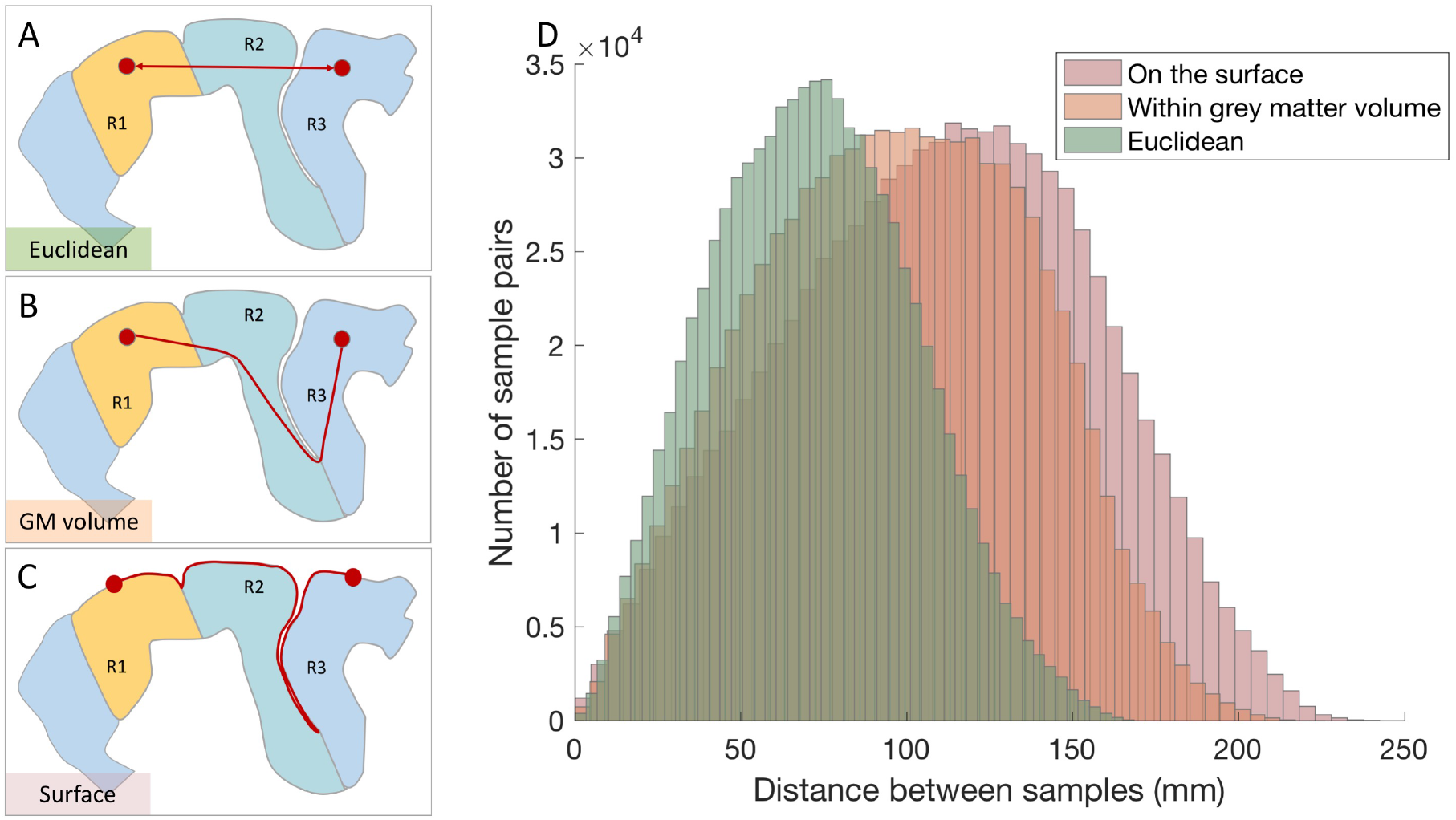
Distances between samples can be evaluated using different methods. Schematic representation of different approaches to calculate the distances between samples. A) Euclidean distance, defined as the shortest distance between two points. B) Distance within grey matter defined as the shortest distance within the grey matter volume. This measure is implemented by representing each voxel in the cortex as a node and creating a three-dimensional network where the shortest distances between brain regions are found using Dijkstra’s algorithm (Dijkstra, 1959). The resulting distances will approximate the distances between regions within the cortical volume. C) Distance on a mesh-based representation of the cortical surface, where gene expression samples are assigned to vertices in the mesh and the shortest distance is calculated as a shortest path between them. Both Euclidean and grey matter distances can be calculated using volumetric parcellation schemes, while estimating the distance on the cortical surface requires generating a surface-based cortical parcellation. D) Distributions for pairwise sample distances calculated using the three approaches. Euclidean distance estimates are lower compared to both distances estimated within grey matter volume and on the surface.

Spatial effects are most easily examined in the context of analyses of correlated gene expression (CGE). Such analyses focus on patterns of pair-wise or multivariate transcriptional coupling between regions, where transcriptional coupling is estimated as a correlation between regional expression profiles. Such measures of CGE can then be related to some inter-regional IDP, such as a measure of functional or structural connectivity (Richiardi et al., 2015, Fulcher and Fornito, 2016, Arnatkevičiūtė et al., 2018). Figure 9A shows that CGE decays sharply as a function of increasing spatial distance (on the pial surface) between regions in the cortex; relationships for other distance measures are qualitatively similar (see Figure S5). In line with previous findings in different species (Fulcher and Fornito, 2016, Arnatkevičiūtė et al., 2018), the dependence of CGE on distance can be approximated as an exponential (Figure 9A) and therefore the residuals of the exponential fit could be further used in the analyses (Figure 9B). Extending this relationship to the whole-brain including samples from both cortex and subcortex is complicated by a strong anti-correlation between cortical and subcortical gene expression (Hawrylycz et al., 2015). Thus, separate normalization procedures for cortical and subcortical regions and corrections for different types of region pairs can be applied (see supplementary material S5 and Figure S6 for more details). Note also that the dependence of CGE on distance can vary as a function of the gene set and parcellation (see Figure S7).

**Fig 9.**
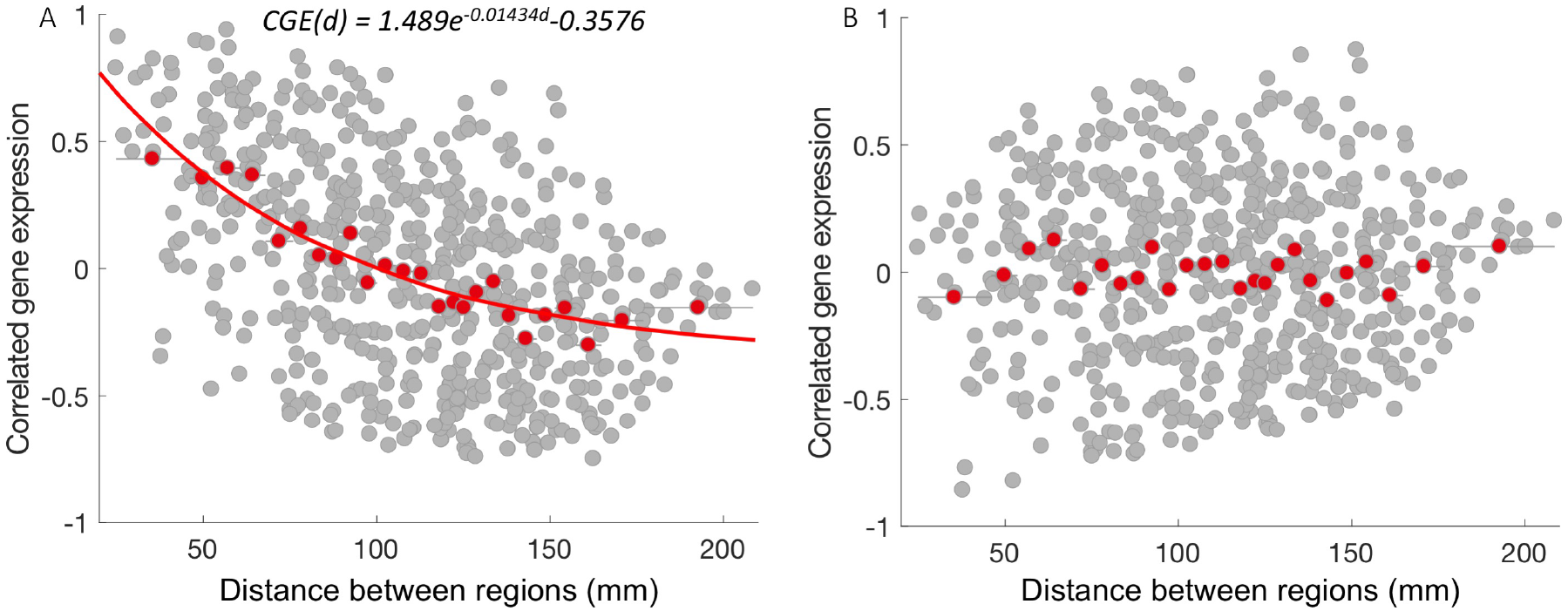
Spatial effect in correlated gene expression. Relationship between correlated gene expression (CGE) and separation distance for the cortical regions in the left hemisphere. A) CGE as a function of separation distance. The red line represents an exponential fit *CGE*(*d*) = 1.498*e*^−0.01434*d*^ − 0.3576. B) CGE residuals after removing the exponential trend; CGE between pairs of regions are represented in grey dots and red dots represent the mean value in 25 equiprobable distance bins after the correction. CGE calculated using all 10,028 genes (after intensity-based filtering and probe selection based on correlation to RNA-seq data).

Characterizing and removing distance dependence can be relatively straightforward in analyses of CGE. Addressing spatial relationships in analyses of regional gene expression can be more challenging since distance is defined between pairs of regions, whereas a regional expression value is a property of a single region. Some promising strategies to deal with this issue involve comparing observed findings relative to an appropriate null model. One class of methods uses spatially constrained permutation of the original data. Arbitrarily-defined regions are not independent form one another, so some spatial constraints are required to account for these dependencies during permutation. As an example, a block permutation algorithm implemented by Vértes et al. (2016) accounted for spatial relationships between regions by aggregating areas into spatially contiguous subsets (blocks) according to the Desikan-Killiany atlas, and then permuting the resulting blocks rather than individual regions. Vasa et al. (2018) introduced a spatial permutation test based on the rotation of regional coordinates in the spherical projection such that the relative spatial relationships between regions are preserved. Matching between original and rotated coordinates, therefore, allows the regional measure to be permuted while controlling for spatial contiguity and hemispheric symmetry. Burt et al. (2017) used a spatial lagged autocorrelation model to characterise the spatial dependency between observed gene expression values. While these approaches provide some valid options, thorough evaluation of these null models is an important avenue of future work.

## Conclusions

Imaging transcriptomics provides an unprecedented opportunity to uncover the molecular basis of large-scale brain organization. Given the rapid development of this field and its heavy reliance on publicly available data, there is a pressing need for standardized data processing pipelines that will facilitate the comparison of findings across studies. Our analysis delineates seven core steps of a basic workflow and demonstrates how choices at each step may affect the final expression measures. We summarize some preliminary recommendations for best practice in Table 4.

**Table 4.**
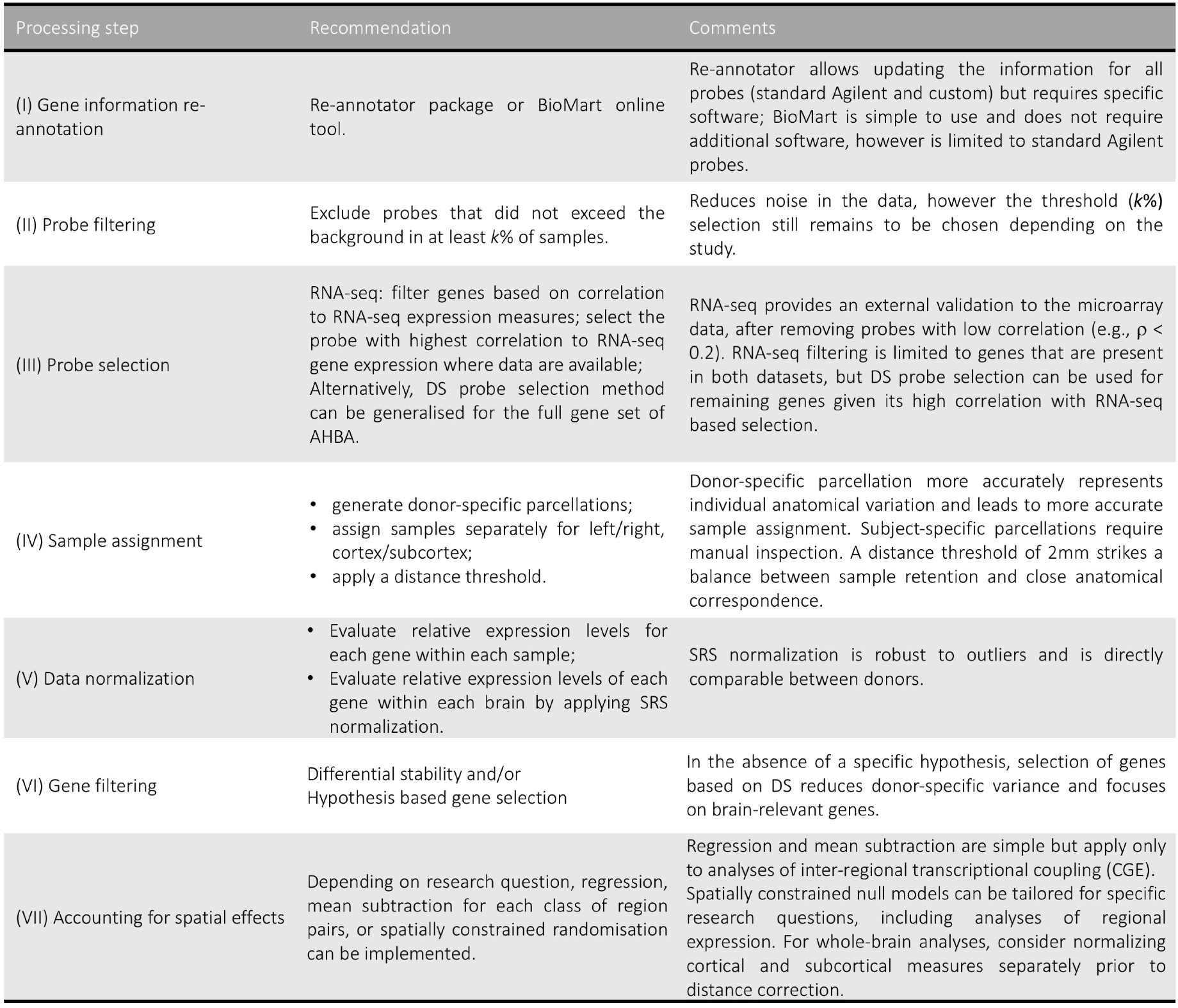
Recommendations and practical considerations for each data processing step.

Considerable further work is required, particularly in the development of methods for addressing spatial correlations in the data. The development of standardized workflows will be essential to ensure reproducibility, particularly as gene expression atlases become more widely available and increase in their sophistication (Lein et al., 2007, Harris et al., 2010, Miller et al., 2014a). We have focused here on the processing of expression measures and removal of inherent biases in the data. Another area requiring further work is the development of appropriate statistical methods for relating IDPs to transcriptomic measures. For example, there is considerable variability in the software packages used for enrichment analyses, each of which makes different assumptions and uses different annotations of genes to gene ontology and other categories (Rhee et al., 2008). It will be important to understand how the available choices for analyzing these data affect reproducibility.

## Acknowledgments

We would like to thank A/Prof David Powell and Dr. Sarah Williams for valuable comments regarding the gene expression data processing. AF was supported by the Australian Research Council (ID: FT130100589) and National Health and Medical Research council (IDs: 1146292). BDF was supported by an NHMRC Early Career Fellowship (ID: 1089718).

## Supporting Information

### supplementary material S1

#### Details on AHBA

The AHBA microarray gene expression data consists of 3702 samples from six neurotypical adult brains. Several hundred samples (mean ± standard deviation: 617 ± 241) were collected from cortical, subcortical, brainstem and cerebellar regions in each brain and profiled for genome-wide gene expression using custom Agilent 8 × 60K cDNA chip which consists of a standard Whole Human Genome Microarray Kit, 4 × 44K (Design ID: 014850) and more than 18,000 custom-generated probes created specifically for AHBA in order to increase the genetic coverage. Originally, 48,171 of all 58,692 probes were annotated to a gene, resulting in a set of 20,787 unique genes with expression measures. In addition to gene expression data, the AHBA provides a binary indicator when the level of a given transcript exceeds background. This assignment is done on the basis of two criteria: 1) a two-sided *t*-statistic comparing the mean signal of a probe to the background (*p* <0.01); and 2) retaining only background-subtracted signals that are 2.6 × above the standard deviation of the background.

Each probe in the AHBA is associated with a numerical ID and a platform-specific label or name. If a probe is assigned to represent a unique gene it is also characterized with a range of gene-specific labels such as gene symbol and an entrez gene ID – a stable identifier for a gene generated by the Entrez Gene database at the National Center for Biotechnology Information (NCBI). If required, probe sequences can be accessed using the Allen Institute’s website application programming interface (see supplementary material S6) while Agilent probe sequences can also be downloaded through the manufacturer’s website (https://earray.chem.agilent.com/earray/). Probe sequences can also be found in the figshare repository https://figshare.com/s/441295fe494375aa0c13. Note that the expression levels of a single gene can be measured using multiple probes that correspond to a different part of the gene sequence. In the AHBA, 93% of all genes are annotated to more than one probe.

Probe-level data are available for each of 3702 tissue samples taken throughout the brain. Different brain regions were sampled across each of the six AHBA donors to maximize spatial coverage. Each tissue sample is associated with a numeric structure ID, name and structure label (‘cortex’, ‘cerebellum’, or ‘brainstem’) in addition to the MRI voxel coordinates in native image space and MNI coordinates in standard space that can be used for matching samples to the stereotaxic space. The AHBA also provides RNA-seq data for a subset of tissue samples in two donor brains (*n* = 120 each). It consists of expression values for more than 22,000 genes presented in fragment count (number of reads matching a given gene) and TMP (Transcripts Per Kilobase Million - normalized read count with regards to read and transcript length) formats. RNA-seq method quantifies the transcription by directly sequencing each molecule in high-throughput manner, therefore providing a more precise measurement of levels of transcripts without the prior knowledge of the DNA sequence of interest (Wang et al., 2009).

Magnetic resonance images—T1-weighted, T2-weighted, T2-weighted gradient echo and FLAIR—were collected prior to dissection of each brain for anatomic visualization. Diffusion tensor images were collected for two brains (H0351_2001 and H0351_2002). Detailed information about image acquisition sequences is presented in the technical white paper (Allen Human Brain Atlas, 2013).

### supplementary material S2

#### Enrichment analysis

Software: version 3.1.2 version of ErmineJ software (Gillis et al., 2010);

Biological process GO annotations: obtained from GEMMA (Zoubarev et al., 2012) as `Generic_human_ncbiIds_noParents.an.txt` downloaded on May 16, 2018.

Gene Ontology terms and definitions: obtained from archive.geneontology.org/latest-termdb/go_daily-termdb.rdf-xml.gz on May 16 2018.

The analyses were performed only on the biological process annotations.

#### Intensity-based filtering

To test if intensity-based filtering (where probes are filtered based on the binary indication of expression levels exceeding the background) targets any specific functional gene groups, we performed the gene score resampling (GSR) analysis. Avoiding the potential bias of overestimating the influence of genes that are represented with multiple probes scores for the GRS analysis were determined at gene (rather than probe) level by: (i) calculating the proportion of samples with expression values exceeding the background using binary indicator provided by AHBA for each probe; (ii) if more than one probe was available for a gene, a probe with the highest proportion of samples exceeding the background was selected to represent that gene. As a result, each of the 20,232 genes was assigned a score indicating the proportion of samples with expression levels exceeding the background. The analysis was performed focusing on genes with low scores in order to determine what functional gene groups are affected by the intensity-based filtering. The mean score in a GO group was selected to summarize it, using full resampling with 10^6^ iterations. FDR-corrected *p*-values (across around 7000 GO categories) were used to summarize the effect. The significant GO categories (at *p* < 0.05) include non brain-specific processes such as sensory perception, chemotaxis, cell killing, and immune response among others (a list of TOP 100 GO categories is presented in supplementary file enrichmentExpression.csv).

#### RNA-seq – microarray non-overlap

The usage of RNA-seq expression measures for probe selection in microarray data is limited to genes that are present in both datasets. Given that out of 20,232 genes in the microarray data ~ 13% (*n* =2623) genes are not present in the RNA-seq dataset, we aimed to test if any brain-specific functional groups of genes are over-represented in this set as these genes would be excluded from the further analysis. If this is the case, then probe selection based on the correlation to RNA-seq data would not be an optimal solution due to the loss of relevant information. On the other hand, if the excluded set of genes does not correspond to brain-specific genes, then the exclusion of those genes may not be a critical issue. Overrepresentation analysis (ORA) was performed for each biological process GO category with 5 to 100 genes available taking the mean score in a GO group to summarize it. FDR-corrected *p*-values (across around 7000 GO categories) were used to summarize the effect. The significant GO categories (at *p* < 0.05) include general processes such as septin assembly and organization as well as the negative regulation of RNA splicing among others (presented in supplementary file enrichmentExpression.csv) and are not related to brain-specific biological processes.

#### Microarray and RNA-seq correlation

In order for RNA-seq gene expression measures to provide a valid reference when selecting a probe, microarray and RNA-seq measures should be at least weakly correlated. Given that for a number of genes the maximum correlation between microarray and RNA-seq expression measures is very low, the probe selection based on such low correlations will be invalid. Therefore, we first exclude probes exhibiting a low correlation (*ρ* < 0.2) to RNA-seq data resulting in the exclusion of 6725 genes. To evaluate the functional groups of genes that were removed, we performed an overrepresentation analysis (ORA). Avoiding the potential bias of overestimating the influence of genes that are represented with multiple probes, scores for the ORA analysis were determined at gene (rather than probe) level by: (i) calculating correlation between microarray and RNA-seq expression for each probe in two subjects; (ii) estimating the mean correlation for each probe across two subjects; (iii) if the maximum correlation value was lower than 0.2, a gene was excluded and assigned an arbitrary value of 0 to serve as a binary indicator of exclusion; otherwise a value of 1 was assigned to represent a gene. As a result, each of 17,609 genes were assigned a score indicating whether it was excluded due to low correlation to RNA-seq data. ORA was performed for each biological process GO category with 5 to 100 genes available taking the mean score in a GO group to summarize it. FDR-corrected *p*-values (across around 7000 GO categories) were used to summarize the effect. The significant (at *p* < 0.05) GO categories include general processes such as immune response, DNA modification and regulation of transposition among others (in Supplementary File enrichmentExpression.csv).

To further verify that genes with higher correlations between microarray and RNA-seq were related to neuronal connectivity and communication related processes we performed gene score resampling analysis (GSR). Avoiding the potential bias of overestimating the influence of genes that are represented with multiple probes scores for the GSR analysis were determined at gene (rather than probe) level by: (i) calculating correlation between microarray and RNA-seq expression for each probe in two subjects; (ii) estimating the mean correlation for each probe across two subjects; (iii) if more than one probe was available for a gene, the maximum correlation value was selected to represent that gene. As a result, each of the 17,609 genes that are present in both microarray and RNA-seq datasets was assigned a score indicating the maximum correlation between microarray and RNA-seq expression values across matching structures.

Focusing on genes with high scores in order to determine which functional gene groups are more likely to be correlated with RNA-seq expression values, we treated larger scores as indicative of the signal. Gene score resampling was performed for each biological process GO category with 5 to 100 genes available taking the mean score in a GO group to summarize it, using full resampling with 10^6^ iterations. FDR-corrected *p*-values (across around 7000 GO categories) were used to summarize the effect. The significant (at *p* < 0.05) GO categories include brain-specific processed such as ensheathment of neurons, oligodendrocyte development, transmission of nerve impulse, glial cell development, central nervous system myelination, synaptic vesicle transport and action potential among others. Of the TOP 100 significant GO categories, around 50% are related to brain-specific processes (presented in Supplementary File enrichmentExpression.csv; brain-related processes are colored in green).

### supplementary material S3

#### Comments on differences between probe selection methods

Two of the variance-based methods—those based on choosing probes with high coefficient of variation or maximum variance—aim to select probes that vary most across the brain. This approach is based on the logic that investigators are often interested in genes that show variation in expression across brain regions. However, probes with higher variance tend to have lower mean intensity because a lower hybridization leads higher signal variability (Quackenbush, 2002). Indeed, we find a negative relationship between expression variance and mean intensity across probes with expression values exceeding the background (average probe intensity > 3; *ρ* = −0.44, *p* < 0.001, Spearman’s rank correlation, see Figure S1).

Choosing a probe based on the highest loading on the first PC aims to select the probe with the most representative expression pattern based on the probe-to-probe variance-covariance matrix. If the probes are not correlated, the first PC will not be representative. Correlations between probes selected using variance-based and consistency/intensity-based approaches are lower than those obtained through random selection. This suggests that variance and intensity/consistency-based approaches favour probes with more different expression measures, meaning that probes with the highest variance would tend on average to be less consistent than randomly selected probes and vice versa (Figure 4A, for the comparison between probe selection methods after QC filtering is applied see Figure S2).

The most popular approach is to summarize gene expression as a mean of all available probes (see Table 2). This method shows high similarity to all other methods. Given that probes can measure different parts of the same gene with different sensitivity, expression measures quantified using different probes are likely not to be equivalent. In this case summarising expression of the gene by calculating the mean of all available probes is likely to reduce this variability. Consistency-based probe selection selects probes with the most consistent regional expression patterns across the six brains in the AHBA, using a measure called differential stability (DS), first introduced in Hawrylycz et al. (2015). The first analysis of the AHBA data indicated that regional variation in expression levels across anatomical structures was strongly correlated between different brains, thus between-region variation in expression should dominate between-subject variance. Choosing a probe with consistent regional variation assumes that this variation is real, and that any between-subject variance is noise. Such an approach is justifiable in analyses where investigation of regional variations in gene expression are the goal.

### supplementary material S4

#### Evaluating distances between samples

All distances between samples were calculated on a set of samples that were mapped onto individual parcellations applying a 2 mm distance threshold as described in Step 4. Voxel coordinates provided by the AHBA that were used to map samples to subject-specific parcellations are derived from images in subject space and could not be combined to estimate distances between samples from different subjects. Therefore, for both Euclidean distances and distances within cortical surfaces we used MNI coordinates provided by the AHBA for each sample to estimate the pairwise distances between them given that MNI coordinates are derived in the same space for all subjects.

**Euclidean distances** were calculated using the function `pdist2` in MATLAB 2016b^1^. Distances between pairs of regions were estimated as an average of distances between samples within them.

**Distances within cortical volume** were calculated using an HCPMMP1 (Glasser et al., 2016) parcellation in the MNI space (downloaded from https://neurovault.org/collections/1549/, file `MMP_in_MNI_corr.nii`) by: i) changing the strides of the image from [–13 –2] to [–123] in order to change the image orientation; ii) rendering the brain parcellation in MNI space as a 3D matrix; ii) converting the original MNI sample coordinates to voxel-based coordinates that correspond to the parcellation loaded in a 3D matrix format; iv) finding the closest coordinate in the parcellation for each sample and mapping a sample to that location; v) rendering the parcellation as a graph with each voxel representing a node; vi) implementing Dijkstra’s algorithm (Dijkstra, 1959) to calculate the closest distances between samples within the GM volume using `shortestpath` function in MATLAB. Distances between pairs of regions were calculated as an average of distances between samples within them.

**Distance on the cortical surface** were calculated by: i) making a separate file for samples of each subject in volumetric space using their MRI voxel coordinates; ii) transforming volumetric files to surface files; iii) creating a separate file for each sample in surface space; iv) mapping samples from the native subject-specific space to fsaverage space; v) choosing a single vertex for each sample by selecting a vertex with maximum value when multiple were available; vi) calculating the distances between samples (mapped to vertices) on the cortical surface using the `toolbox_fast_marching` toolbox in MATLAB. Distances between pairs of regions were calculated as an average of distances between samples within them.

### supplementary material S5

#### Spatial relationship between CGE and distance

Figure S6B shows the relationship between CGE and separation distance for three different types of regional pairs: (i) intra-cortical region pairs, which show relatively high CGE at all distances (blue); (ii) intra-subcortical region pairs, which show a linearly decreasing relationship at short distances (red); and (iii) cortical-subcortical region pairs, which demonstrate mostly negative CGE at all distances (yellow). This variability precludes simple removal of a global trend as in (Fulcher and Fornito, 2016) and suggests a separate correction should be applied for each class of connection. Figure S6C shows the result of removing the mean CGE at each equiprobable distance bin separately for each connection class. Even after this correction, intracortical CGE values appear to be underestimated while cortico-subcortical are overestimated. Another method for addressing the inherent differences between cortical and subcortical gene expression profiles is to normalize the expression data separately for these two anatomical divisions [e.g., Anderson et al. (2018)]. With this approach, expression values are scaled relative to other values within just the cortex, or within just the subcortex. Division-specific normalization allows a gene to score highly if its expression in the subcortex is high relative to other subcortical regions, even if its expression relative to the cortex may be low (and vice-versa), at the expense of distorting the magnitude relationships between cortical and subcortical values. With this approach, the negative correlation between cortical and subcortical regions is reduced with some region pairs demonstrating relatively similar gene expression profiles (Figure S6D). As presented in Figure S6E, the relationship between CGE and separation distance is now more qualitatively similar for all three groups and can be corrected using simple mean subtraction across distance bins (Figure S6F).

### supplementary material S6

**API: probe sequences** for the first 10,000 rows.

http://api.brain-map.org/api/v2/data/query.xml?criteria=model::Probe,rma::criteria,products[id$eq2],rma::include,gene,predicted_sequence,rma::options[only$eq%27probes.name,probes.type,probes.ncbi_accession_number,probes.gi,genes.entrez_id,genes.acronym,sequences.sequence_length,sequences.sequence_data%27],[num_rows$eq10000][start_row$eq0]

**Fig S1.**
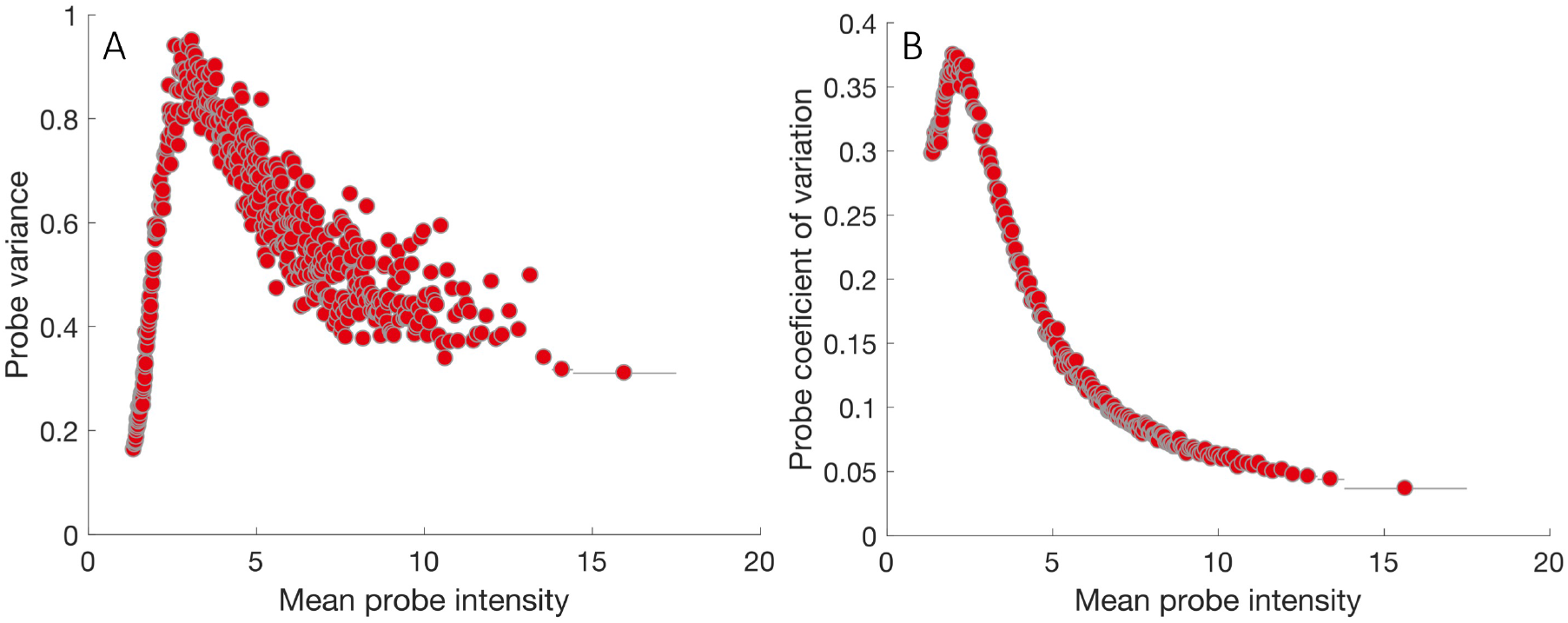
The relationship between average probe intensity, variance and coefficient of variation. The relationship between average probe intensity and A) variance, and B) coefficient of variation in 500 and 250 equiprobable intensity bins respectively, shown as a circle (bin centers) and a horizontal line (bin extent). The relationship between probe intensity and variance is positive for low intensity probes (intensity< 3); for higher intensity probes increasing intensity results in decreasing variance. The same trend is evident for the relationship between coefficient of variation and intensity.

**Fig S2.**
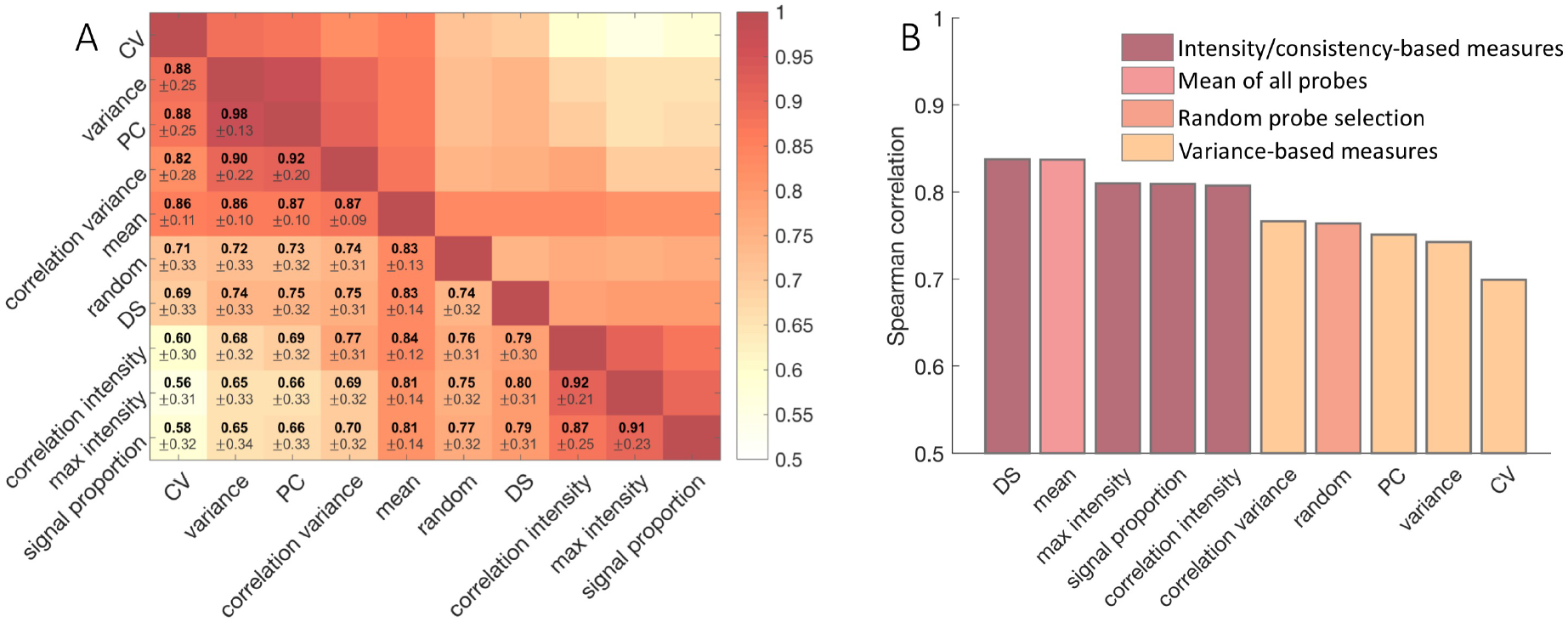
Average correlation between summary expression scores for genes annotated to multiple probes, where a single representative probe is chosen based on different criteria after intensity-based filtering. A) Average correlation between summary expression scores for genes annotated to multiple probes, where a single representative probe is chosen based on different criteria: CV, variance, PC, signal proportion, DS, correlation variance, correlation intensity, mean (see Table 2) or selecting a representative probe at random (correlation values averaged over 100 runs). The average correlation is computed over 11,190 genes with multiple probe annotations after intensity-based filtering. B) Average correlation between probes selected using RNA-seq expression (by selecting the probe that is correlated to the RNA-seq data the most) and other methods (ordered by decreasing values, based on 7950 genes that (i) were present in both microarray and RNA-seq datasets; (ii) were correlated to RNA-seq (*ρ* > 0.2, Spearman rank correlation) to ensure that RNA-seq based probe selection provides a meaningful estimate; (iii) had more than one probe available after intensity-based filtering).

**Fig S3.**
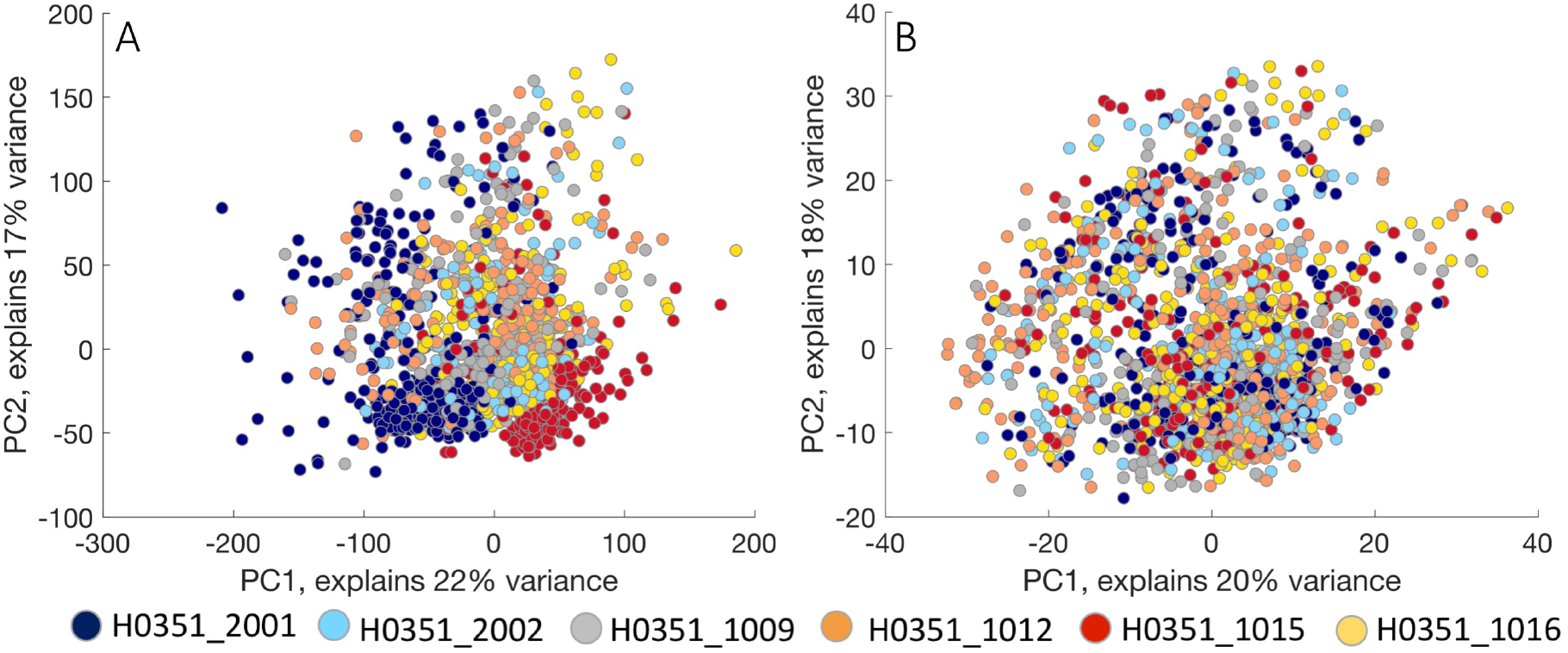
Inter-individual differences in gene expression and the effect of normalization in the whole brain. A) Original gene expression data for cortical and subcortical samples in principal component space. Data from different donors are represented in different colours. Samples from different subjects occupy different parts of the low-dimensional gene expression space. Cortical samples (right in A) are slightly separated from subcortical samples (left in A). Panel B represent gene expression data in principal component space normalized using scaled robust sigmoid (SRS) demonstrating much more clear separation between cortical and subcortical samples. After normalization samples no longer segregate by donor.

**Fig S4.**
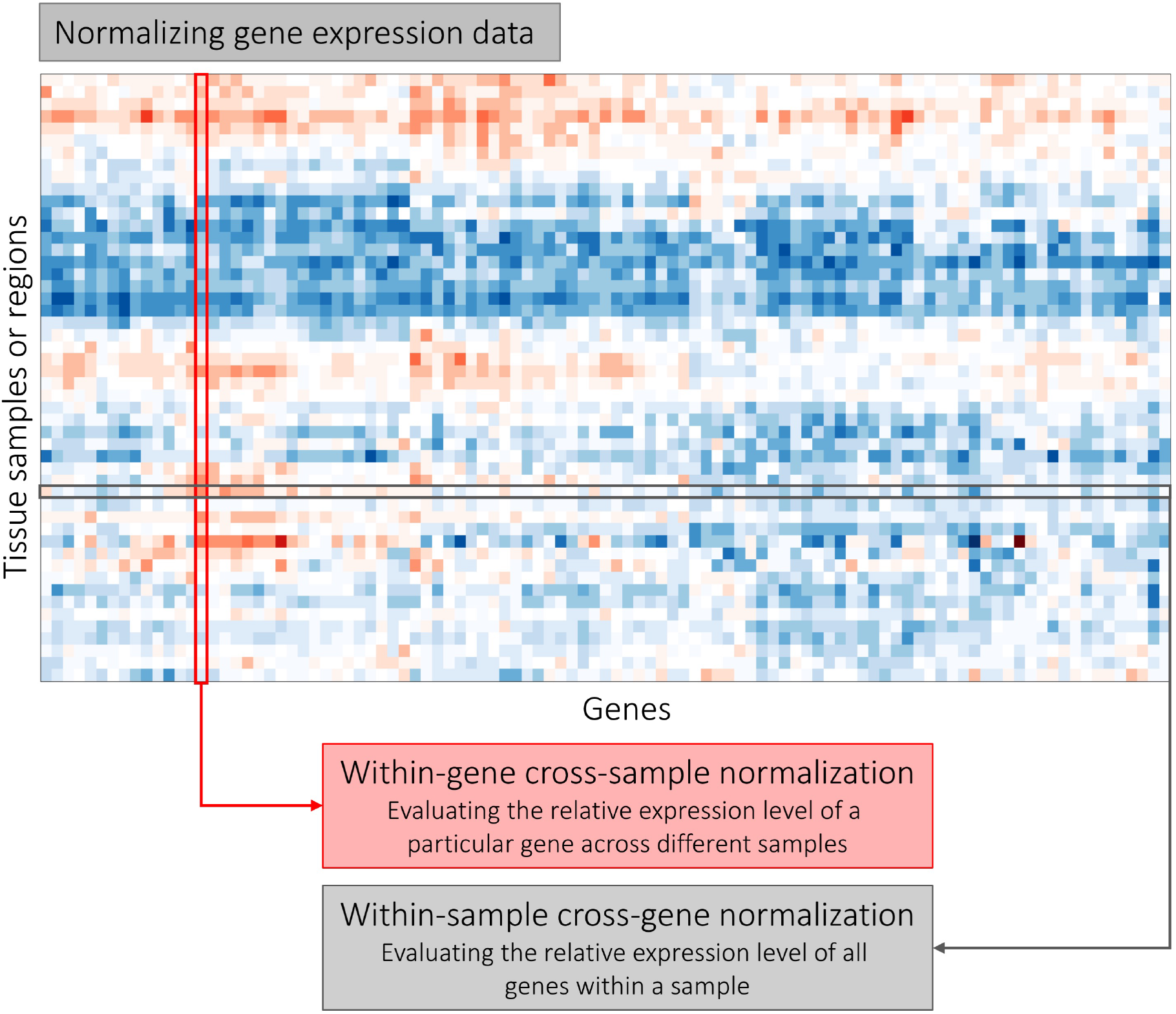
Gene expression normalization. The schematic representation of cross-gene and cross-sample normalization. Within-sample cross-gene normalization estimates the relative expression level of all available genes within a given sample (grey row). Within-gene cross sample normalization estimates the relative expression of a particular gene across all available samples (red column).

**Fig S5.**
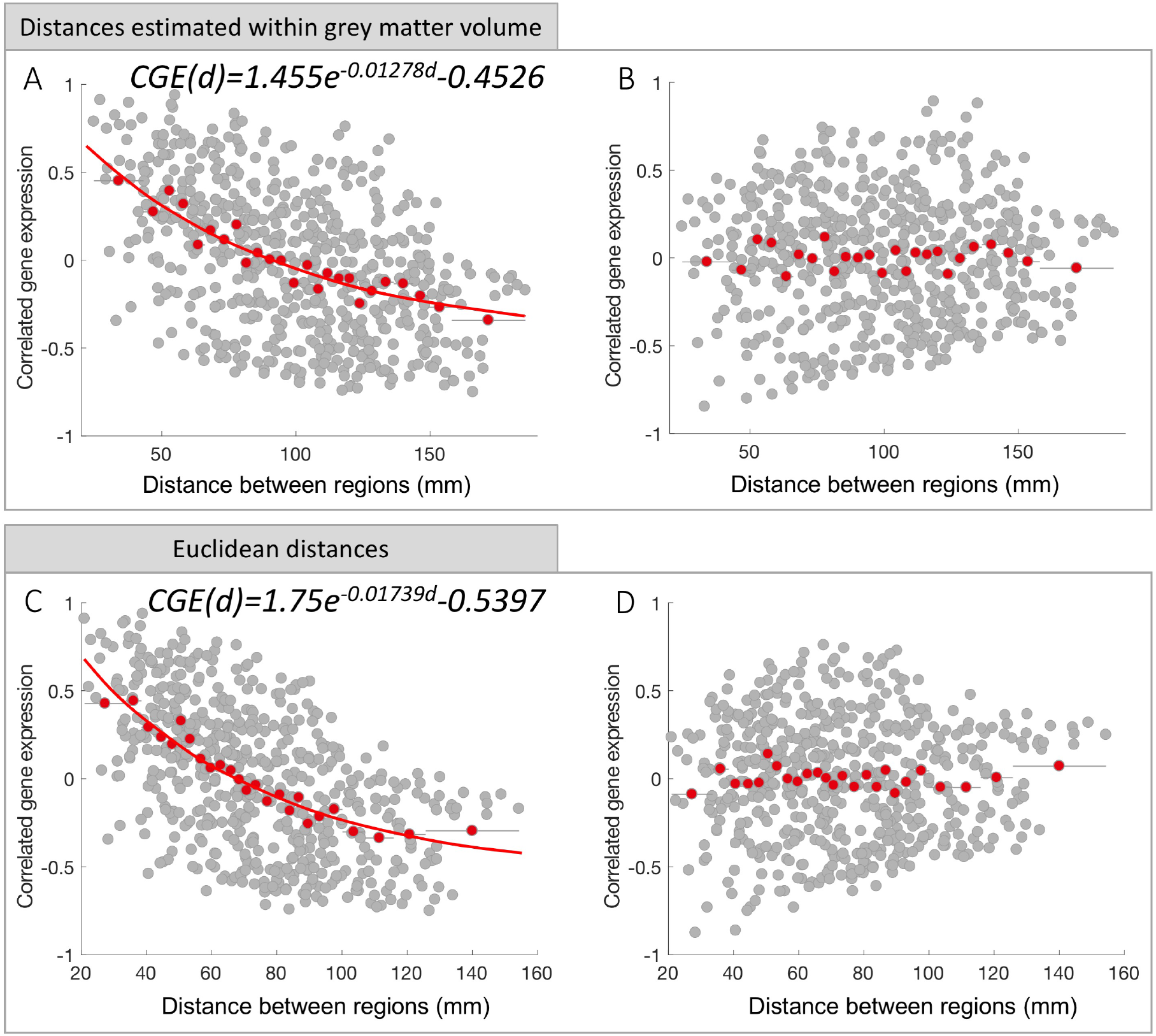
The relationship between CGE and inter-regional distance estimated within grey matter volume and as Euclidean distance. Top: the relationship between CGE and distance when the distance between regions calculated within grey matter volume. Bottom: the relationship between CGE and distance when the distance between regions calculated as Euclidean distance. A,C) CGE as a function of separation distance where CGE between pairs of regions are represented in grey dots and red dots represent the mean value in 25 equiprobable distance bins; The red line represents an exponential fit. A) *CGE*(*d*) = 1.455*e*^−0.01278*d*^ − 0.4526; C) *CGE*(*d*) = 1.75*e*^−0.01739*d*^ − 0.5397. B,D) Residuals after removing the exponential trend in each case; CGE calculated using all 10,028 genes (after intensity-based filtering and probe selection based on correlation to RNA-seq data).

**Fig S6.**
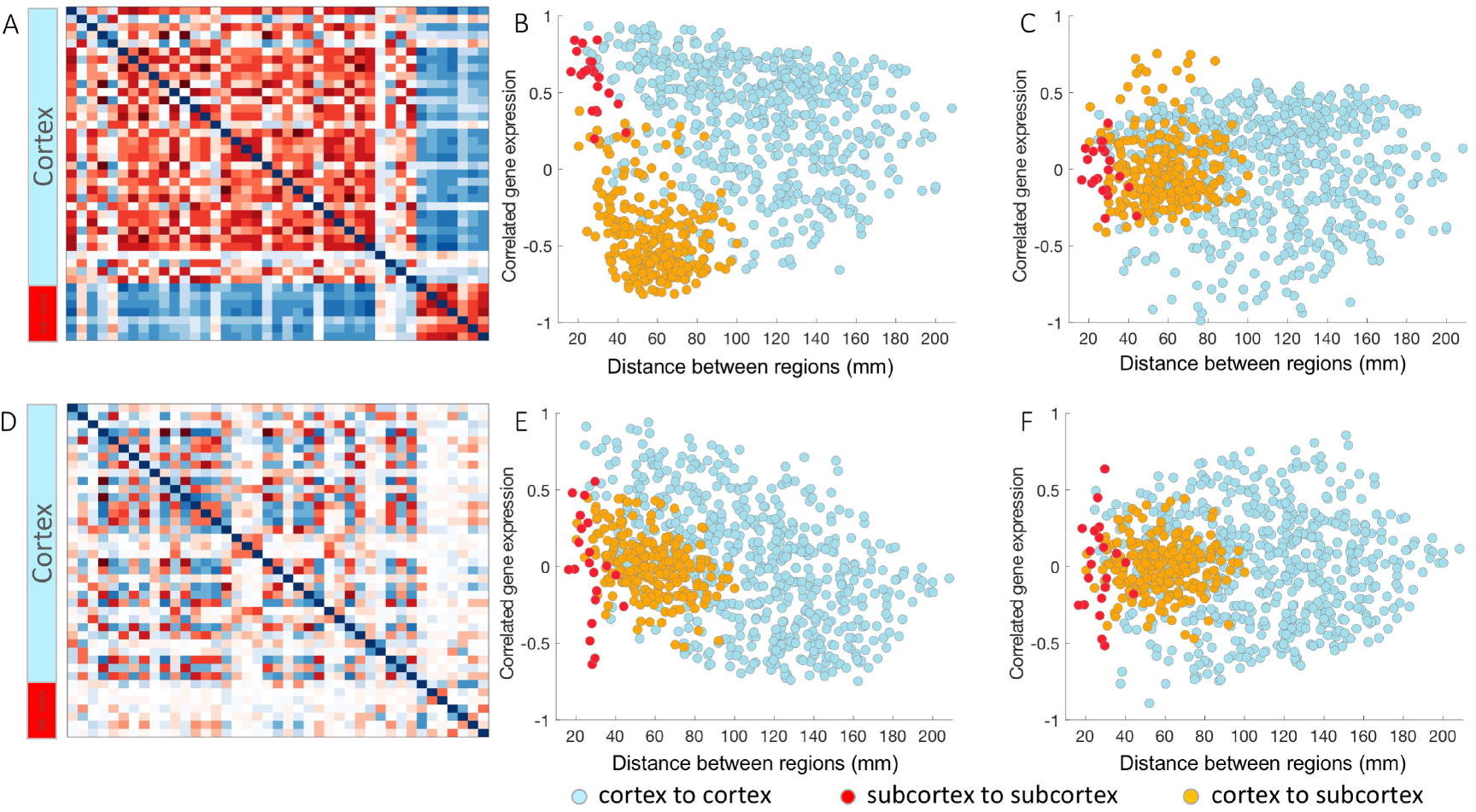
The relationship between CGE and inter-regional distance for cortical and subcortical regions. Top row: A) Matrix of CGE values for the left hemisphere, including both cortical and subcortical regions, in which expression values have been normalized for both cortical and subcortical regions together; B) CGE as a function of separation distance where CGE between different subsets of regions are represented in different colours: within cortex — light blue, within subcortex — red, between cortex and subcortex — yellow; C) CGE residuals after removing the spatial effect for each subset of regions separately by subtracting the average of each bin, where different colours represent subsets of connections as above. D) Matrix of CGE values for the left hemisphere, including both cortical and subcortical regions, in which expression values have been normalized separately for cortical and subcortical regions; E) CGE as a function of separation distance, where normalization has been performed on cortical and subcortical regions separately. Different colours represent subsets of connections as above; F) CGE residuals after removing the spatial effect for each subset of regions separately (when normalization performed on cortical and subcortical regions separately) by subtracting the average of each bin where different colours represent subsets of connections as above. Distances between cortical regions were evaluated on the cortical surface while distances between subcortical regions as well as between cortical and subcortical regions estimated as Euclidean.

**Fig S7.**
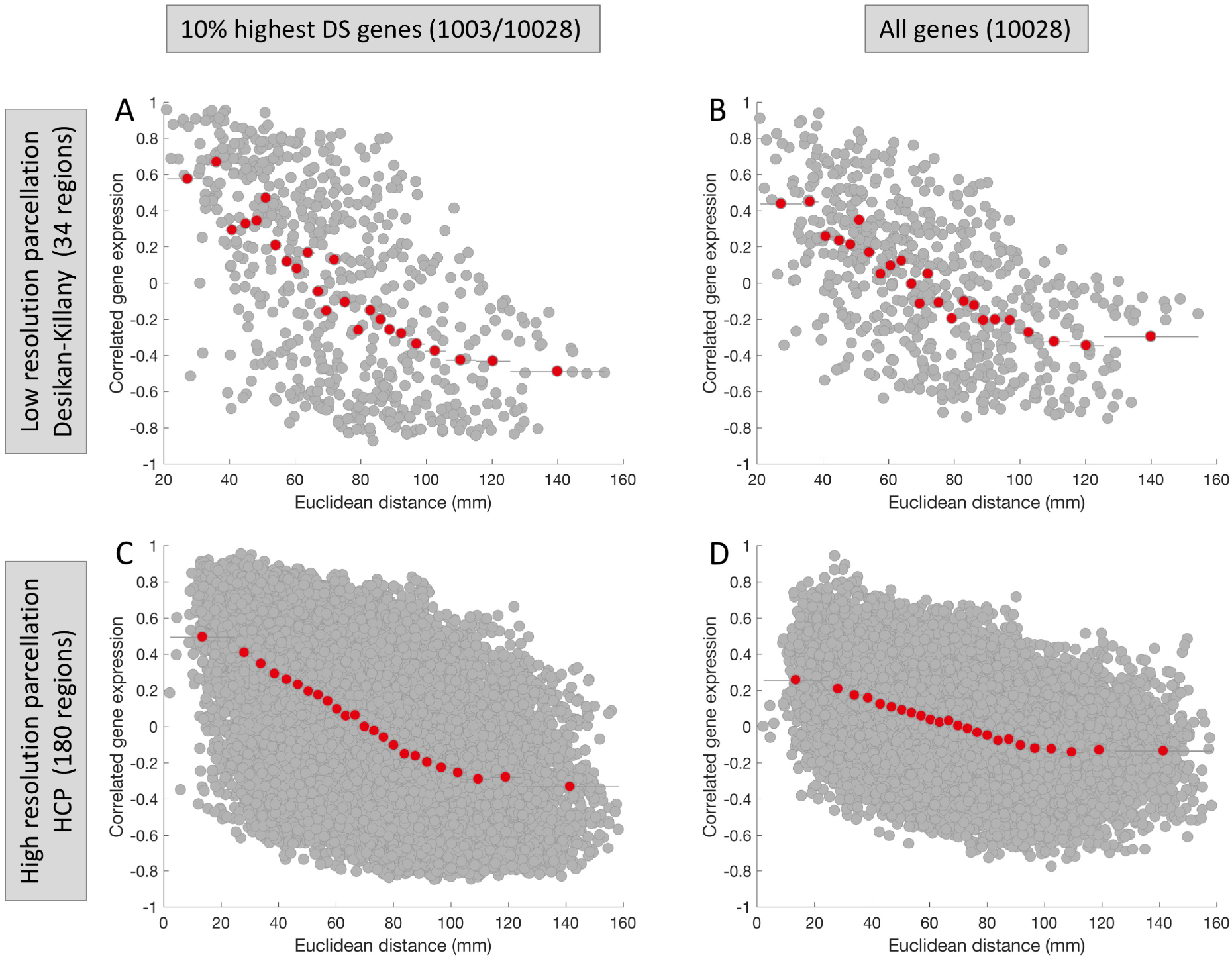
The relationship between CGE and inter-regional distance for high and low resolution parcellations using a full set of genes and a set of only highest DS genes. The relationship between CGE and inter-regional distance for different resolution cortical parcellations and different sets of genes. Top row: low resolution Desikan-Killany parcellation (Desikan et al., 2006) (34 regions). Bottom row: high resolution HCPMMP1 (Glasser et al., 2016) parcellation (180 regions). First column: CGE calculated using only 10% of highest DS genes that are most consistently expressed across subjects and regions. Second column: CGE calculated using all 10,028 genes. The relationship between CGE and inter-regional distance also depends on the subset of genes chosen for the calculation, such that high DS genes show a stronger association with inter-regional distance. This effect is more pronounced in the higher resolution parcellation.

## Footnotes

1 MATLAB is a product of Mathworks

## References

Acevedo-Triana, C.A., León, L.A., Cardenas, F.P., Cardenas, F.P., 2017. Comparing the expression of genes related to serotonin (5-HT) in C57BL/6J mice and humans based on data available at the Allen Mouse Brain Atlas and Allen Human Brain Atlas. Neurol. Res. Int. 2017, 1–14. doi:10.1155/2017/7138926.

Allen Human Brain Atlas, 2013. AHBA documentation. URL: http://help.brain-map.org/display/humanbrain/Documentation.

Anderson, K.M., Krienen, F.M., Choi, E.Y., Reinen, J.M., Yeo, B.T.T., Holmes, A.J., 2018. Gene expression links functional networks across cortex and striatum. Nat. Commun. 9, 1428. doi:10.1038/s41467-018-03811-x.

Arloth, J., Bader, D.M., Röh, S., Altmann, A., 2015. Re-Annotator: Annotation Pipeline for Microarray Probe Sequences. PLoS One 10, e0139516. doi:10.1371/journal.pone.0139516.

Arnatkevičiūtė, A., Fulcher, B.D., Pocock, R., Fornito, A., 2018. Hub connectivity, neuronal diversity, and gene expression in the Caenorhabditis elegans connectome. PLoS Comput. Biol. 14, e1005989. doi:10.1371/journal.pcbi.1005989.

Arslan, A., 2015. Genes, brains, and behavior: imaging genetics for neuropsychiatric disorders. J. Neuropsychiatry Clin. Neurosci. 27, 81–92. doi:10.1176/appi.neuropsych.13080185.

Bakken, T.E., Miller, J.A., Ding, S.L., Sunkin, S.M., Smith, K.A., et al., 2016. A comprehensive transcriptional map of primate brain development. Nature 535, 367–375. doi:10.1038/nature18637.

Berchtold, N.C., Cribbs, D.H., Coleman, P.D., Rogers, J., Head, E., Kim, R., Beach, T., Miller, C., Troncoso, J., Trojanowski, J.Q., Zielke, H.R., Cotman, C.W., 2008. Gene expression changes in the course of normal brain aging are sexually dimorphic. Proc. Natl. Acad. Sci. U. S. A. 105, 15605–10. doi:10.1073/pnas.0806883105.

Birdsill, A.C., Walker, D.G., Lue, L., Sue, L.I., Beach, T.G., 2011. Postmortem interval effect on RNA and gene expression in human brain tissue. Cell Tissue Bank. 12, 311–318. doi:10.1007/s10561-010-9210-8.

Burt, J.B., Demirtas, M., Eckner, W.J., Navejar, N.M., Ji, J.L., Martin, W.J., Bernacchia, A., Anticevic, A., Murray, J.D., 2017. Hierarchy of transcriptomic specialization across human cortex captured by myelin map topography. bioRxiv, 199703 doi:10.1101/199703.

Chen, L., Chu, C., Zhang, Y.H., Zhu, C., Kong, X., Huang, T., Cai, Y.D., 2016. Analysis of gene expression profiles in the human brain stem, cerebellum and cerebral cortex. PLoS One 11, e0159395. doi:10.1371/journal.pone.0159395.

Choi, J.K., Kim, S.C., 2007. Environmental effects on gene expression phenotype have regional biases in the human genome. Genetics 175, 1607–13. doi:10.1534/genetics.106.069047.

Cioli, C., Abdi, H., Beaton, D., Burnod, Y., Mesmoudi, S., 2014. Differences in human cortical gene expression match the temporal properties of large-scale functional networks. PLoS One 9, e115913. doi:10.1371/journal.pone.0115913.

Ciric, R., Wolf, D.H., Power, J.D., Roalf, D.R., Baum, G.L., Ruparel, K., Shinohara, R.T., Elliott, M.A., Eickhoff, S.B., Davatzikos, C., Gur, R.C., Gur, R.E., Bassett, D.S., Satterthwaite, T.D., 2017. Benchmarking of participant-level confound regression strategies for the control of motion artifact in studies of functional connectivity. Neuroimage 154, 174–187. doi:10.1016/J.NEUROIMAGE.2017.03.020.

Colantuoni, C., Lipska, B.K., Ye, T., Hyde, T.M., Tao, R., Leek, J.T., Colantuoni, E.A., Elkahloun, A.G., Herman, M.M., Weinberger, D.R., Kleinman, J.E., 2011. Temporal dynamics and genetic control of transcription in the human prefrontal cortex. Nature 478, 519–23. doi:10.1038/nature10524.

Cole, S.W., 2009. Social regulation of human gene expression. Curr. Dir. Psychol. Sci. 18, 132–137. doi:10.1111/j.1467-8721.2009.01623.x.

Desikan, R.S., Ségonne, F., Fischl, B., Quinn, B.T., Dickerson, B.C., Blacker, D., Buckner, R.L., Dale, A.M., Maguire, R.P., Hyman, B.T., Albert, M.S., Killiany, R.J., 2006. An automated labeling system for subdividing the human cerebral cortex on MRI scans into gyral based regions of interest. Neuroimage 31, 968–980. doi:10.1016/j.neuroimage.2006.01.021.

Dijkstra, E.W., 1959. A note on two problems in connexion with graphs. Numer. Math. 1, 269–271. doi:10.1007/BF01386390.

Eising, E., Huisman, S.M., Mahfouz, A., Vijfhuizen, L.S., Anttila, V., et al., 2016. Gene co-expression analysis identifies brain regions and cell types involved in migraine pathophysiology: a GWAS-based study using the Allen Human Brain Atlas. Hum. Genet. 135, 425–439. doi:10.1007/s00439-016-1638-x.

Fare, T.L., Coffey, E.M., Dai, H., He, Y.D., Kessler, D.A., Kilian, K.A., Koch, J.E., LeProust, E., Marton, M.J., Meyer, M.R., Stoughton, R.B., Tokiwa, G.Y., Wang, Y., 2003. Effects of atmospheric ozone on microarray data quality. Anal. Chem. 75, 4672–5. URL: http://www.ncbi.nlm.nih.gov/pubmed/14632079.

Ferrari, R., Hernandez, D.G., Nalls, M.A., Rohrer, J.D., Ramasamy, A., et al., 2014. Frontotemporal dementia and its subtypes: a genome-wide association study. Lancet Neurol. 13, 686–699. doi:10.1016/S1474-4422(14)70065-1.

Fertuzinhos, S., Li, M., Kawasawa, Y., Ivic, V., Franjic, D., Singh, D., Crair, M., Šestan, N., 2014. Laminar and temporal expression dynamics of coding and noncoding RNAs in the mouse neocortex. Cell Rep. 6, 938–950. doi:10.1016/j.celrep.2014.01.036.

Forest, M., Iturria-Medina, Y., Goldman, J.S., Kleinman, C.L., Lovato, A., Oros Klein, K., Evans, A., Ciampi, A., Labbe, A., Greenwood, C.M., 2017. Gene networks show associations with seed region connectivity. Hum. Brain Mapp. doi:10.1002/hbm.23579.

Fraser, H.B., Khaitovich, P., Plotkin, J.B., Pääbo, S., Eisen, M.B., 2005. Aging and gene expression in the primate brain. PLoS Biol. 3, e274. doi:10.1371/journal.pbio.0030274.

French, L., Paus, T., 2015. A FreeSurfer view of the cortical transcriptome generated from the Allen Human Brain Atlas. Front. Neurosci. 9, 323. doi:10.3389/fnins.2015.00323.

Fulcher, B.D., Fornito, A., 2016. A transcriptional signature of hub connectivity in the mouse connectome. Proc. Natl. Acad. Sci. 113, 1513302113. doi:10.1073/pnas.1513302113,

Fulcher, B.D., Little, M.A., Jones, N.S., 2013. Highly comparative time-series analysis: the empirical structure of time series and their methods. J. R. Soc. Interface 10, 20130048. doi:10.1098/rsif.2013.0048.

Futcher, B., Latter, G.I., Monardo, P., McLaughlin, C.S., Garrels, J.I., 1999. A sampling of the yeast proteome. Mol. Cell. Biol. 19, 7357–68. URL: http://www.ncbi.nlm.nih.gov/pubmed/10523624.

Gillis, J., Mistry, M., Pavlidis, P., 2010. Gene function analysis in complex data sets using ErmineJ. Nat. Protoc. 5, 1148–59. doi:10.1038/nprot.2010.78.

Glass, D., Viñuela, A., Davies, M.N., Ramasamy, A., Parts, L., et al., 2013. Gene expression changes with age in skin, adipose tissue, blood and brain. Genome Biol. 14, R75. doi:10.1186/gb-2013-14-7-r75.

Glasser, M.F., Coalson, T.S., Robinson, E.C., Hacker, C.D., Harwell, J., Yacoub, E., Ugurbil, K., Andersson, J., Beckmann, C.F., Jenkinson, M., Smith, S.M., Van Essen, D.C., 2016. A multi-modal parcellation of human cerebral cortex. Nature 536, 171–178. doi:10.1038/nature18933.

Goel, P., Kuceyeski, A., LoCastro, E., Raj, A., 2014. Spatial patterns of genome-wide expression profiles reflect anatomic and fiber connectivity architecture of healthy human brain. Hum. Brain Mapp. 35, 4204–4218. doi:10.1002/hbm.22471.

Gorgolewski, K.J., Varoquaux, G., Rivera, G., Schwarz, Y., Ghosh, S.S., Maumet, C., Sochat, V.V., Nichols, T.E., Poldrack, R.A., Poline, J.B., Yarkoni, T., Margulies, D.S., 2015. NeuroVault.org: a web-based repository for collecting and sharing unthresholded statistical maps of the human brain. Front. Neuroinform. 9, 8. doi:10.3389/fninf.2015.00008.

Goyal, M.S., Hawrylycz, M., Miller, J.A., Snyder, A.Z., Raichle, M.E., 2014. Aerobic glycolysis in the human brain is associated with development and neotenous gene expression. Cell Metab. 19, 49–57. doi:10.1016/j.cmet.2013.11.020.

Greenbaum, D., Colangelo, C., Williams, K., Gerstein, M., 2003. Comparing protein abundance and mRNA expression levels on a genomic scale. Genome Biol. 4, 117. doi:10.1186/gb-2003-4-9-117.

Gryglewski, G., Seiger, R., James, G.M., Godbersen, G.M., Komorowski, A., Unterholzner, J., Michenthaler, P., Hahn, A., Wadsak, W., Mitterhauser, M., Kasper, S., Lanzenberger, R., 2018. Spatial analysis and high resolution mapping of the human whole-brain transcriptome for integrative analysis in neuroimaging. Neuroimage 176, 259–267. doi:10.1016/j.neuroimage.2018.04.068.

Gygi, S.P., Rochon, Y., Franza, B.R., Aebersold, R., 1999. Correlation between protein and mRNA abundance in yeast. Mol. Cell. Biol. 19, 1720–30. URL: http://www.ncbi.nlm.nih.gov/pubmed/10022859.

Harris, T.W., Antoshechkin, I., Bieri, T., Blasiar, D., Chan, J., et al., 2010. WormBase: a comprehensive resource for nematode research. Nucleic Acids Res. 38, D463–D467. doi:10.1093/nar/gkp952.

Hashimoto, R., Ohi, K., Yamamori, H., Yasuda, Y., Fujimoto, M., Umeda-Yano, S., Watanabe, Y., Fukunaga, M., Takeda, M., 2015. Imaging genetics and psychiatric disorders. Curr. Mol. Med. 15, 168–75. doi:10.2174/1566524015666150303104159.

Hawrylycz, M., Miller, J.A., Menon, V., Feng, D., Dolbeare, T., et al., 2015. Canonical genetic signatures of the adult human brain. Nat. Neurosci. 18, 1832–1844. doi:10.1038/nn.4171.

Hawrylycz, M.J., Lein, E.S., Guillozet-Bongaarts, A.L., Shen, E.H., Ng, L., et al., 2012. An anatomically comprehensive atlas of the adult human brain transcriptome. Nature 489, 391–9. doi:10.1038/nature11405.

Hecker, N., Seemann, S.E., Silahtaroglu, A., Ruzzo, W.L., Gorodkin, J., 2017. Associating transcription factors and conserved RNA structures with gene regulation in the human brain. Sci. Rep. 7, 1–16. doi:10.1038/s41598-017-06200-4.

Höglinger, G.U., Melhem, N.M., Dickson, D.W., Sleiman, P.M.A., Wang, L.S., et al., 2011. Identification of common variants influencing risk of the tauopathy progressive supranuclear palsy. Nat. Genet. 43, 699–705. doi:10.1038/ng.859.

Jaksik, R., Iwanaszko, M., Rzeszowska-Wolny, J., Kimmel, M., 2015. Microarray experiments and factors which affect their reliability. Biol. Direct 10, 46. doi:10.1186/s13062-015-0077-2.

Johnson, M.B., Kawasawa, Y.I., Mason, C.E., Krsnik, Ž., Coppola, G., Bogdanović, D., Geschwind, D.H., Mane, S.M., State, M.W., Šestan, N., 2009. Functional and evolutionary insights into human brain development through global transcriptome analysis. Neuron 62, 494–509. doi:10.1016/j.neuron.2009.03.027.

Kang, H.J., Kawasawa, Y.I., Cheng, F., Zhu, Y., Xu, X., et al., 2011. Spatio-temporal transcriptome of the human brain. Nature 478, 483–9. doi:10.1038/nature10523.

Keil, J.M., Qalieh, A., Kwan, K.Y., 2018. Brain transcriptome databases: a user’s guide. J. Neurosci. 10, 1930–17. doi:10.1523/JNEUROSCI.1930-17.2018.

Keo, A., Aziz, N.A., Dzyubachyk, O., van der Grond, J., van Roon-Mom, W.M.C., Lelieveldt, B.P.F., Reinders, M.J.T., Mahfouz, A., 2017. Co-expression patterns between ATN1 and ATXN2 coincide with brain regions affected in Huntington’s disease. Front. Mol. Neurosci. 10, 1–13. doi:10.3389/fnmol.2017.00399.

Kirsch, L., Chechik, G., 2016. On expression patterns and developmental origin of human brain regions. PLoS Comput. Biol. 12, 1–25. doi:10.1371/journal.pcbi.1005064.

Komorowski, A., James, G.M., Philippe, C., Gryglewski, G., Bauer, A., et al., 2016. Association of protein distribution and gene expression revealed by PET and post-mortem quantification in the serotonergic system of the human brain. Cereb. Cortex 27, 117–130. doi:10.1093/cercor/bhw355.

Kouri, N., Ross, O.A., Dombroski, B., Younkin, C.S., Serie, D.J., et al., 2015. Genome-wide association study of corticobasal degeneration identifies risk variants shared with progressive supranuclear palsy. Nat. Commun. 6, 7247. doi:10.1038/ncomms8247.

Krebs, J.E., Lewin, B., Kilpatrick, S.T., Goldstein, E.S., 2014. Lewin’s genes XI. 11th ed. / ed., Jones & Bartlett Learning, Burlington Mass. URL: http://www.worldcat.org/title/lewins-genes-xi/oclc/794228027.

Krienen, F.M., Yeo, B.T.T., Ge, T., Buckner, R.L., Sherwood, C.C., 2016. Transcriptional profiles of supragranular-enriched genes associate with corticocortical network architecture in the human brain. Proc. Natl. Acad. Sci. 113, E469–78. doi:10.1073/pnas.1510903113.

Kukurba, K.R., Montgomery, S.B., 2015. RNA Sequencing and Analysis. Cold Spring Harb. Protoc. 2015, 951–69. doi:10.1101/pdb.top084970.

Kumar, A., Gibbs, J.R., Beilina, A., Dillman, A., Kumaran, R., Trabzuni, D., Ryten, M., Walker, R., Smith, C., Traynor, B.J., Hardy, J., Singleton, A.B., Cookson, M.R., 2013. Age-associated changes in gene expression in human brain and isolated neurons. Neurobiol. Aging 34, 1199–209. doi:10.1016/j.neurobiolaging.2012.10.021.

Lee, A.G., Hagenauer, M., Absher, D., Morrison, K.E., Bale, T.L., Myers, R.M., Watson, S.J., Akil, H., Schatzberg, A.F., Lyons, D.M., 2017. Stress amplifies sex differences in primate prefrontal profiles of gene expression. Biol. Sex Differ. 8, 36. doi:10.1186/s13293-017-0157-3.

Lein, E.S., Hawrylycz, M.J., Ao, N., Ayres, M., Bensinger, A., et al., 2007. Genome-wide atlas of gene expression in the adult mouse brain. Nature 445, 168–176. doi:10.1038/nature05453.

Liu, Z., Rolls, E.T., Zhang, J., Yang, M., Du, J., Gong, W., Cheng, W., Wang, H., Ugurbil, K., Feng, J., 2017. The functional and genetic associations of neuroimaging data: a toolbox. bioRxiv, 178640 doi:10.1101/178640.

Margineantu, D.H., Emerson, C.B., Diaz, D., Hockenbery, D.M., 2007. Hsp90 inhibition decreases mitochondrial protein turnover. PLoS One 2, e1066. doi:10.1371/journal.pone.0001066.

McColgan, P., Gregory, S., Seunarine, K.K., Razi, A., Papoutsi, M., et al., 2018. Brain regions showing white matter loss in Huntington’s disease are enriched for synaptic and metabolic genes. Biol. Psychiatry 83, 456–465. doi:10.1016/J.BIOPSYCH.2017.10.019.

Mexal, S., Berger, R., Adams, C., Ross, R., Freedman, R., Leonard, S., 2006. Brain pH has a significant impact on human postmortem hippocampal gene expression profiles. Brain Res. 1106, 1–11. doi:10.1016/j.brainres.2006.05.043.

Meyer-Lindenberg, A., Weinberger, D.R., 2006. Intermediate phenotypes and genetic mechanisms of psychiatric disorders. Nat. Rev. Neurosci. 7, 818–827. doi:10.1038/nrn1993.

Miller, J.A., Cai, C., Langfelder, P., Geschwind, D.H., Kurian, S.M., Salomon, D.R., Horvath, S., 2011. Strategies for aggregating gene expression data: The collapseRows R function. BMC Bioinformatics 12, 322. doi:10.1186/1471-2105-12-322.

Miller, J.A., Ding, S.L., Sunkin, S.M., Smith, K.A., Ng, L., et al., 2014a. Transcriptional landscape of the prenatal human brain. Nature 508, 199–206. doi:10.1038/nature13185.

Miller, J.A., Menon, V., Goldy, J., Kaykas, A., Lee, C.K., Smith, K.A., Shen, E.H., Phillips, J.W., Lein, E.S., Hawrylycz, M.J., 2014b. Improving reliability and absolute quantification of human brain microarray data by filtering and scaling probes using RNA-Seq. BMC Genomics 15, 154. doi:10.1186/1471-2164-15-154.

Muñoz, K.E., Hyde, L.W., Hariri, A.R., 2009. Imaging genetics. J. Am. Acad. Child Adolesc. Psychiatry 48, 356–61. doi:10.1097/CHI.0b013e31819aad07.

Myers, E.M., Bartlett, C.W., Machiraju, R., Bohland, J.W., 2015. An integrative analysis of regional gene expression profiles in the human brain. Methods 73, 54–70. doi:10.1016/j.ymeth.2014.12.010.

Naumova, O.Y., Palejev, D., Vlasova, N.V., Lee, M., Rychkov, S.Y., Babich, O.N., M. Vaccarino, F., Grigorenko, E.L., 2012. Age-related changes of gene expression in the neocortex: Preliminary data on RNA-Seq of the transcriptome in three functionally distinct cortical areas. Dev. Psychopathol. 24, 1427–1442. doi:10.1017/S0954579412000818.

Negi, S.K., Guda, C., 2017. Global gene expression profiling of healthy human brain and its application in studying neurological disorders. Sci. Rep. 7, 897. doi:10.1038/s41598-017-00952-9.

O’Leary, N.A., Wright, M.W., Brister, J.R., Ciufo, S., Haddad, D., et al., 2016. Reference sequence (RefSeq) database at NCBI: Current status, taxonomic expansion, and functional annotation. Nucleic Acids Res. 44, D733–D745. doi:10.1093/nar/gkv1189.

Pantazatos, S.P., Li, X., 2017. Commentary: BRAIN NETWORKS. Correlated gene expression supports synchronous activity in brain networks. Science 348, 1241–4. Front. Neurosci. 11, 412. doi:10.3389/fnins.2017.00412.

Parkes, L., Fulcher, B., Yücel, M., Fornito, A., 2018. An evaluation of the efficacy, reliability, and sensitivity of motion correction strategies for resting-state functional MRI. Neuroimage 171, 415–436. doi:10.1016/J.NEUROIMAGE.2017.12.073.

Parkes, L., Fulcher, B.D., Yücel, M., Fornito, A., 2017. Transcriptional signatures of connectomic subregions of the human striatum. Genes, Brain Behav. doi:10.1111/gbb.12386.

Power, J.D., Plitt, M., Laumann, T.O., Martin, A., 2017. Sources and implications of whole-brain fMRI signals in humans. Neuroimage 146, 609–625. doi:10.1016/j.neuroimage.2016.09.038.

Power, J.D., Schlaggar, B.L., Petersen, S.E., 2015. Recent progress and outstanding issues in motion correction in resting state fMRI. Neuroimage 105, 536–551. doi:10.1016/J.NEUROIMAGE.2014.10.044.

Quackenbush, J., 2002. Microarray data normalization and transformation. Nat. Genet. 32, 496–501. doi:10.1038/ng1032.

Rhee, S. Y, Wood, V Dolinski, K Draghici, S 2008. Use and misuse of the gene ontology annotations. Nat. Rev. Genet. 9, 509–515. doi:10.1038/ng1032.

Richiardi, J., Altmann, A., Greicius, M., 2017. Distance is not everything in imaging genomics of functional networks: reply to a commentary on correlated gene expression supports synchronous activity in brain networks. bioRxiv doi:https://doi.org/10.1101/132746.

Richiardi, J., Altmann, A., Milazzo, A.C., Chang, C., Chakravarty, M.M.,et al., 2015. Correlated gene expression supports synchronous activity in brain networks. Science 348, 1241–1244. doi:10.1126/science.1255905, arXiv:arXiv:1011.1669v3.

Ripke, S., Neale, B.M., Corvin, A., Walters, J.T.R., Farh, K.H., et al., 2014. Biological insights from 108 schizophrenia-associated genetic loci. Nature 511, 421–427. doi:10.1038/nature13595.

Ritchie, M.E., Phipson, B., Wu, D., Hu, Y., Law, C.W., Shi, W., Smyth, G.K., 2015. limma powers differential expression analyses for RNA-sequencing and microarray studies. Nucleic Acids Res. 43, e47–e47. doi:10.1093/nar/gkv007.

Rittman, T., Rubinov, M., Vértes, P.E., Patel, A.X., Ginestet, C.E., Ghosh, B.C., Barker, R.A., Spillantini, M.G., Bullmore, E.T., Rowe, J.B., 2016. Regional expression of the MAPT gene is associated with loss of hubs in brain networks and cognitive impairment in Parkinson disease and progressive supranuclear palsy. Neurobiol. Aging 48, 153–160. doi:10.1016/j.neurobiolaging.2016.09.001.

Rittman, T., 2017. Maybrain software package. URL: https://github.com/rittman/maybrain.

Rizzo, G., Veronese, M., Expert, P., Turkheimer, F.E., Bertoldo, A., 2016. MENGA: a new comprehensive tool for the integration of neuroimaging data and the Allen Human Brain Transcriptome Atlas. PLoS One 11, e0148744. doi:10.1371/journal.pone.0148744.

Romero-Garcia, R., Warrier, V., Bullmore, E.T., Baron-Cohen, S., Bethlehem, R.A.I., 2018a. Synaptic and transcriptionally downregulated genes are associated with cortical thickness differences in autism. Mol. Psychiatry, 1 doi:10.1038/s41380-018-0023-7.

Romero-Garcia, R., Whitaker, K.J., Váša, F., Seidlitz, J., Shinn, M., Fonagy, P., Dolan, R.J., Jones, P.B., Goodyer, I.M., Bullmore, E.T., Vértes, P.E., 2018b. Structural covariance networks are coupled to expression of genes enriched in supragranular layers of the human cortex. Neuroimage 171, 256–267. doi:10.1016/J.NEUROIMAGE.2017.12.060.

Romme, I.A.C., de Reus, M.A., Ophoff, R.A., Kahn, R.S., van den Heuvel, M.P., 2017. Connectome disconnectivity and cortical gene expression in patients with schizophrenia. Biol. Psychiatry 81, 495–502. doi:10.1016/j.biopsych.2016.07.012.

Satake, W., Nakabayashi, Y., Mizuta, I., Hirota, Y., Ito, C., et al., 2009. Genome-wide association study identifies common variants at four loci as genetic risk factors for Parkinson’s disease. Nat. Genet. 41, 1303–1307. doi:10.1038/ng.485.

Scherer, A., 2009. Batch effects and noise in microarray experiments : sources and solutions. J. Wiley. URL: https://www.wiley.com/en-us/Batch+Effects+and+Noise+in+Microarray+Experiments{%}3A+Sources+and+Solutions-p-9780470741382.

Schwanhäusser, B., Busse, D., Li, N., Dittmar, G., Schuchhardt, J., Wolf, J., Chen, W., Selbach, M., 2013. Corrigendum: Global quantification of mammalian gene expression control. Nature 495, 126–127. doi:10.1038/nature11848.

Shin, J., French, L., Xu, T., Leonard, G., Perron, M., Pike, G.B., Richer, L., Veillette, S., Pausova, Z., Paus, T., 2017. Cell-specific gene-expression profiles and cortical thickness in the human brain. Cereb. Cortex, 1–11 doi:10.1093/cercor/bhx197.

Simón-Sánchez, J., Schulte, C., Bras, J.M., Sharma, M., Gibbs, J.R., et al., 2009. Genome-wide association study reveals genetic risk underlying Parkinson’s disease. Nat. Genet. 41, 1308–1312. doi:10.1038/ng.487.

Spielman, R.S., Bastone, L.A., Burdick, J.T., Morley, M., Ewens, W.J., Cheung, V.G., 2007. Common genetic variants account for differences in gene expression among ethnic groups. Nat. Genet. 39, 226–31. doi:10.1038/ng1955.

Steger, D., Berry, D., Haider, S., Horn, M., Wagner, M., Stocker, R., Loy, A., 2011. Systematic spatial bias in DNA microarray hybridization is caused by probe spot position-dependent variability in lateral diffusion. PLoS One 6, e23727. doi:10.1371/journal.pone.0023727.

Szymański, M., Barciszewski, J., 2002. Beyond the proteome: non-coding regulatory RNAs. Genome Biol. 3, reviews0005. URL: http://www.ncbi.nlm.nih.gov/pubmed/12049667.

Tan, P.P.C., French, L., Pavlidis, P., 2013. Neuron-enriched gene expression patterns are regionally anti-correlated with oligodendrocyte-enriched patterns in the adult mouse and human brain. Front. Neurosci. 7, 5. doi:10.3389/fnins.2013.00005.

Tarca, A.L., Romero, R., Draghici, S., 2006. Analysis of microarray experiments of gene expression profiling. Am. J. Obstet. Gynecol. 195, 373–88. doi:10.1016/j.ajog.2006.07.001.

Thompson, P.M., Stein, J.L., Medland, S.E., Hibar, D.P., Vasquez, A.A., et al., 2014. The ENIGMA Consortium: large-scale collaborative analyses of neuroimaging and genetic data. Brain Imaging Behav. 8, 153–82., doi:10.1007/s11682-013-9269-5.

Trabzuni, D., Ramasamy, A., Imran, S., Walker, R., Smith, C., Weale, M.E., Hardy, J., Ryten, M., Consortium, N.A.B.E., 2013. Widespread sex differences in gene expression and splicing in the adult human brain. Nat. Commun. 4, 2771. doi:10.1038/ncomms3771.

Váša, F., Seidlitz, J., Romero-Garcia, R., Whitaker, K.J., Rosenthal, G., et al. 2018. Adolescent tuning of association cortex in human structural brain networks. Cereb. Cortex 28, 281–294. doi:10.1093/cercor/bhx249.

Vértes, P.E., Rittman, T., Whitaker, K.J., Romero-Garcia, R., Váša, F., Kitzbichler, M.G., Wagstyl, K., Fonagy, P., Dolan, R.J., Jones, P.B., Goodyer, I.M., Bullmore, E.T., 2016. Gene transcription profiles associated with inter-modular hubs and connection distance in human functional magnetic resonance imaging networks. Philos. Trans. R. Soc. B Biol. Sci. 371, 735–769. doi:10.1098/rstb.2015.0362.

Wang, Z., Gerstein, M., Snyder, M., 2009. RNA-Seq: a revolutionary tool for transcriptomics. Nat. Rev. Genet. 10, 57–63. doi:10.1038/nrg2484.

Whitaker, K.J., Vértes, P.E., Romero-Garcia, R., Váša, F., Moutoussis, M., et al., 2016. Adolescence is associated with genomically patterned consolidation of the hubs of the human brain connectome. Proc. Natl. Acad. Sci. U. S. A. 113, 9105–10. doi:10.1073/pnas.1601745113.

Yokoyama, J.S., Karch, C.M., Fan, C.C., Bonham, L.W., Kouri, N., et al., 2017. Shared genetic risk between corticobasal degeneration, progressive supranuclear palsy, and frontotemporal dementia. Acta Neuropathol. 133, 825–837. doi:10.1007/s00401-017-1693-y.

Zhu, Y., Wang, L., Yin, Y., Yang, E., 2017. Systematic analysis of gene expression patterns associated with postmortem interval in human tissues. Sci. Rep. 7, 5435. doi:10.1038/s41598-017-05882-0.

Zoubarev, A., Hamer, K.M., Keshav, K.D., Luke Mccarthy, E., Santos, J.R.C., Van rossum, T., Mcdonald, C., Hall, A., Wan, X., Lim, R., Gillis, J., Pavlidis, P., 2012. Gemma: A resource for the reuse, sharing and meta-analysis of expression profiling data. Bioinformatics 28, 2272–2273. doi:10.1093/bioinformatics/bts430.

